# Opposing lineage specifiers induce a pro-tumor hybrid-identity state in lung adenocarcinoma

**DOI:** 10.1101/2024.12.02.626384

**Authors:** Gabriela Fort, Henry Arnold, Soledad Camolotto, Rushmeen Tariq, Anna Waters, Kayla O’Toole, Eric L. Snyder

## Abstract

The ability of cancer cells to alter their identity, known as lineage plasticity, is crucial for tumor progression and therapy resistance. In lung adenocarcinoma (LUAD), tumor progression is characterized by a gradual loss of lineage fidelity and the emergence of non-pulmonary identity programs. This can lead to hybrid-identity (hybrid-ID) states in which developmentally incompatible identity programs are co-activated within individual cells. However, the molecular mechanisms underlying these identity shifts remain incompletely understood. Here, we identify the gastrointestinal (GI) transcriptional regulator HNF4α as a critical driver of tumor growth and proliferation in KRAS-driven LUAD. In LUAD cells that express the lung lineage specifier NKX2-1, HNF4α can induce a GI/liver-like state by directly binding and activating its canonical targets. HNF4α also forms an aberrant protein complex with NKX2-1, which disrupts NKX2-1 localization and dampens pulmonary identity within hybrid-ID LUAD. Sustained signaling through the RAS/MEK pathway is critical for maintaining the hybrid-ID state. Moreover, RAS/MEK inhibition augments NKX2-1 chromatin binding at pulmonary-specific genes and induces resistance-associated pulmonary signatures. Finally, we demonstrate that HNF4α depletion enhances sensitivity to pharmacologic KRAS^G12D^ inhibition. Collectively, our data show that co-expression of opposing lineage specifiers leads to a hybrid identity state that can drive tumor progression and dictate response to targeted therapy in LUAD.

## INTRODUCTION

Changes in cancer cell identity, also known as lineage plasticity, drive tumor progression and therapy resistance, presenting a substantial obstacle to cancer treatment. Malignant progression of lung adenocarcinoma (LUAD), the leading cause of cancer-related deaths globally, is driven not solely by genetic alterations, but also by epigenetic changes that confer enhanced plasticity, permitting tumor cells to adopt disparate differentiation states during tumor evolution (LaFave et al., 2020; Marjanovic et al., 2020; Tavernari et al., 2021; Yang et al., 2022). Targeted therapies have revolutionized the treatment landscape for LUAD, but rapid onset of resistance driven by both genetic and epigenetic mechanisms limits the efficacy of these therapies (Araujo et al., 2024; Awad et al., 2021; Canon et al., 2019; Li et al., 2024; Tanaka et al., 2021). Defining the precise molecular mechanisms underlying identity changes in LUAD could unveil novel therapeutic strategies that target identity-specific vulnerabilities or restrain LUAD to a less aggressive differentiation state.

Changes in LUAD cell identity can occur via switch-like or graded mechanisms. Our group found that loss of the master pulmonary lineage-specifying transcription factor NKX2-1, or TTF1, triggers a switch-like change from an alveolar to a gastric differentiation state (Snyder et al., 2013). Mechanistically, this transition is mediated by the pioneer factors FoxA1 and FoxA2 (FoxA1/2), which relocalize from pulmonary to gastric targets following NKX2-1 deletion and impart epigenetic modifications at gastric marker genes (Camolotto et al., 2018; Gillis et al., 2023; Snyder et al., 2013). The majority of human LUAD (~75%) retains expression of NKX2-1 and undergoes more gradual, graded changes in differentiation state during natural progression. In the *Kras/Trp53* mutant (KP) genetically-engineered mouse model (GEMM), NKX2-1-positive LUAD cells gradually transition from a state that is transcriptionally and epigenetically similar to their predominant cell of origin (the alveolar-type 2 (AT2) cell) and adopt disparate differentiation states over time marked by expression of genes associated with other pulmonary or gastrointestinal (GI) cell types (LaFave et al., 2020; Marjanovic et al., 2020; Yang et al., 2022). Progression through these intermediate identity states is postulated to be critical for LUAD cells to reach a poorly differentiated (EMT-like) state with high metastatic potential (Winslow et al., 2011).

One intermediate state in the KP model, which has been annotated as “gastric-like” (Marjanovic et al., 2020), exhibits co-expression of pulmonary and GI transcripts culminating in a “hybrid-identity” (hybrid-ID) state in which multiple differentiation programs are simultaneously activated. This is exemplified by single-cell RNA-sequencing (scRNA-seq) and protein-level data illustrating co-expression of NKX2-1 and HNF4α, a master regulator of identity in the normal GI tract and liver that also robustly marks the gastric-like state (Marjanovic et al., 2020; Orstad et al., 2022). This state is transcriptionally distinct from the high-plasticity cell state (HPCS), and both emerge at a similar timepoint during KP tumor progression (Marjanovic et al., 2020). We recently showed that the pioneer factors FoxA1/2 are critical drivers of growth and acquisition of hybrid-ID differentiation states in NKX2-1-positive LUAD (Orstad et al., 2022). Mechanistically, FoxA1/2 activate aspects of pulmonary differentiation by modulating cell-type-specific binding of NKX2-1, which contributes to growth inhibition following ablation of FoxA1/2 (Orstad et al., 2022). FoxA1/2 are also required for expression of GI-like identity programs, including endodermal lineage-defining transcription factors and marker genes such as HNF4α. However, it is unknown whether activation of GI programs is an additional mechanism by which FoxA1/2 promote tumor growth, and whether HNF4α is a functional driver, or simply a marker, of these identity states in NKX2-1-positive LUAD. Importantly, hybrid-ID states marked by co-expression of NKX2-1 and HNF4α have been identified in human LUAD (Kawai et al., 2024; Kriegsmann et al., 2018; Sugano et al., 2013).

Here we leverage complementary murine and human models to show that HNF4α is a key driver of growth and GI identity programs in NKX2-1-positive LUAD. We also describe an unexpected role for HNF4α in dampening pulmonary differentiation, likely through direct interactions with NKX2-1, which inhibits NKX2-1 binding to canonical pulmonary targets. Finally, we show that RAS/MEK signaling maintains a hybrid-ID state in NKX2-1-positive LUAD by altering chromatin binding of NKX2-1 and define HNF4α as a driver of intrinsic resistance to KRAS inhibition.

## RESULTS

### HNF4α drives growth in NKX2-1-positive LUAD

To interrogate the importance of HNF4α for NKX2-1-positive LUAD tumor growth, we first queried PanCancer LUAD data from The Cancer Genome Atlas (TCGA) database. After filtering for *NKX2-1*-high tumors and excluding biologically distinct mucinous tumors based on pathology reports, we found that high expression of *HNF4A* or its target *EPS8L3* confers a worse prognosis (overall survival) compared to patients with lower *HNF4A* (or *EPS8L3)* expression (**Figure 1A and S1A**). We observed no differences in the distribution of most baseline characteristics and demographics between the *HNF4A*-high and *HNF4A*-low groups (including sex, race, age, and overall stage) (**Supplemental Table S1**).

**Figure 1.**
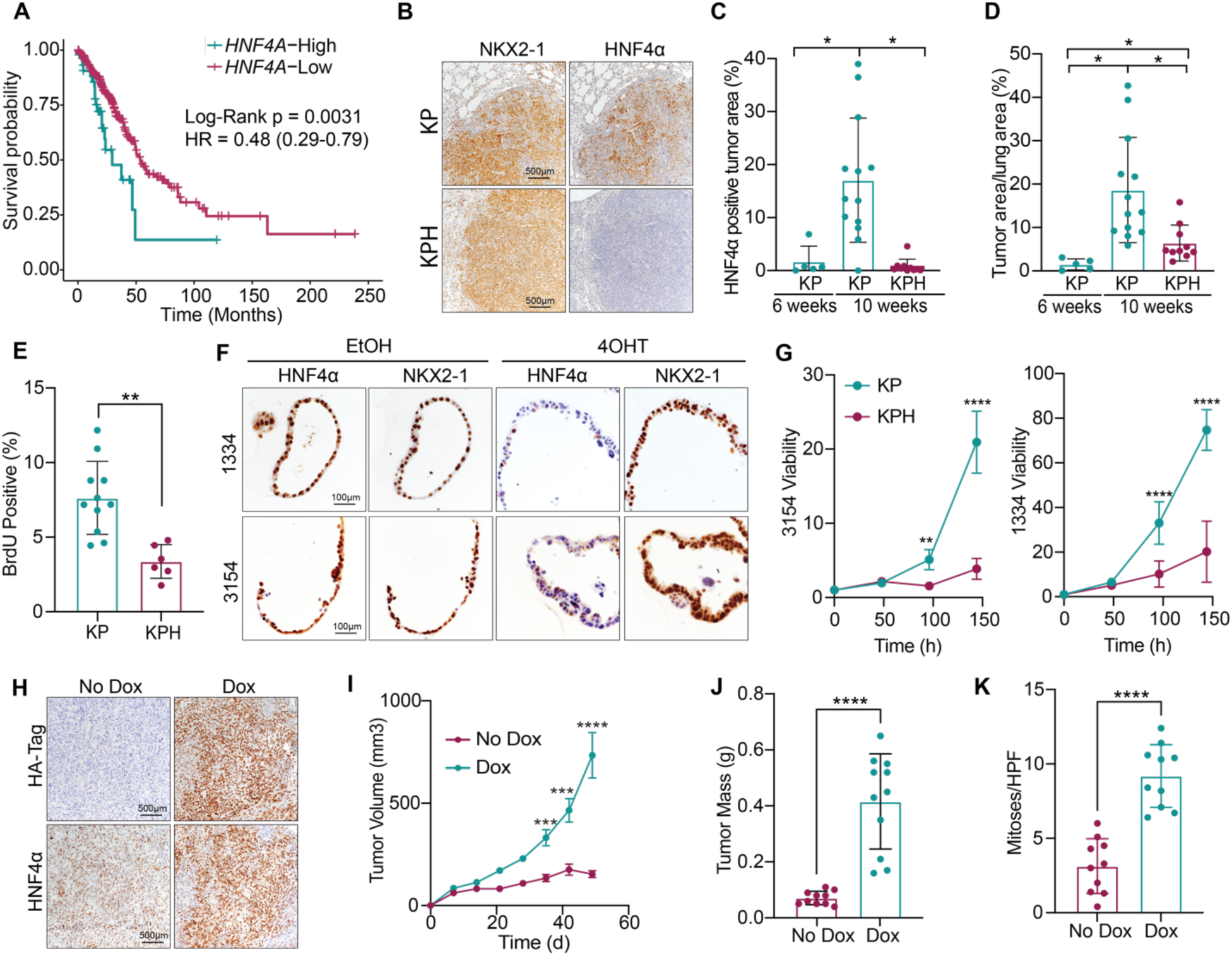
HNF4α promotes the growth of NKX2-1-positive LUAD. **A.** Kaplan-Meier curve of PanCancer LUAD TCGA samples stratified based on high expression of *NKX2-1* and high (n=47) or low (n=339) expression of *HNF4A*, excluding mucinous tumors. Log-Rank test and CoxPH hazard ratio with 95% CI shown. **B.** Representative IHC images of grade 3 tumors from KP/KPH tumors demonstrating protein co-expression of NKX2-1 and HNF4α in subsets of tumor cells. Scale bar = 500μM. **C-E.** Mice were intubated with adenoviral mCMV-FlpO and lungs were harvested at either 6 weeks (n=5) post-tumor initiation or treated with tamoxifen (n=10) or vehicle (n=15) and harvested 4 weeks later. **C.** Quantification of HNF4α-positive tumor area compared to normal lung area (unpaired t tests; *p=0.0117,***p=0.0003; error bars, SD). **D.** Quantification of overall tumor burden (tumor area/total lung area) (unpaired t tests; *p=0.024,**p<=0.007; error bars, SD). **E.** BrdU quantification from IHC staining of grade 3 tumors (unpaired t test; **p=0.001; error bars, SD). **F.** Representative IHC images of DP organoids derived from KPH mice, treated with vehicle (EtOH) or 4-hydroxytamoxifen (4OHT) to delete *Hnf4a*. Scale bar = 100μM. **G.** Presto Blue viability assay on KP (control) or KPH (*Hnf4a*-deleted) DP organoid lines (one representative replicate shown of n=3 biological replicates; unpaired t tests at two timepoints, ***p=0.0002, ****p<0.0001; error bars, SD). **H-K.** H1651 cells were transduced with a dox-inducible, HA-tagged HNF4α overexpression lentivector and subcutaneously implanted into the flanks of NSG mice fed normal chow (n=10) or dox-containing chow (n=11). **H.** Representative IHC images from outgrown subcutaneous tumors demonstrating HNF4α protein overexpression. Scale bar = 500μM. **I.** Tumor volume over time (n=2 biological replicates; unpaired t test day 35 p=0.0002, day 42 p=0.0002, day 49 p<0.0001; error bars, SEM). **J.** Tumor mass at study endpoint (n=2 biological replicates; unpaired t test ****p<0.0001; error bars, SD). **K.** Quantification of mitoses per high-power field (HPF) (n=2 biological replicates; unpaired t test ****p<0.0001; error bars, SD).

HNF4α expression increases during LUAD progression from lower to higher grade tumors and marks a plastic, gastric-like, hybrid-ID state in LUAD (LaFave et al., 2020; Marjanovic et al., 2020; Yang et al., 2022) (**Figure 1B-C**). However, it is unknown whether HNF4α expression drives tumor growth or merely correlates with higher grade disease. To functionally interrogate the role of HNF4α in tumor growth, we generated a sequential recombination GEMM enabling temporally controlled disruption of *Hnf4a* expression in established KRAS-driven lung tumors in vivo. Administration of FlpO recombinase to the lung epithelium in this model induces expression of oncogenic KRAS^G12D^ (Young et al., 2011), deletion of the tumor suppressor *Trp53* (Lee et al., 2012), and *Cre^ERT2^* expression from the ubiquitously active *Rosa26* locus (Schonhuber et al., 2014). Tamoxifen administration then induces *Hnf4a* deletion in tumor cells (Hayhurst et al., 2001) (**Figure S1B**). We utilized *Kras^FSF-G12D^;p53^Frt/Frt^;Rosa26^FSF-CreERT2^;Hnf4a^F/F^* (KPH) mice to delete *Hnf4a* in established tumors via tamoxifen injection at 6 weeks-post tumor initiation and profiled tumors 4 weeks later (10 weeks post tumor initiation). Notably, control (KP) tumors contained a subset of tumor cells harboring high expression of both NKX2-1 and HNF4α (**Figure 1B**). Tamoxifen administration effectively ablated protein expression of HNF4α in KPH tumor cells (**Figure 1B-C**). *Hnf4a* deletion also significantly impaired tumor growth over 4 weeks (**Figure 1D**) and decreased tumor cell proliferation (**Figure 1E**), indicating that HNF4α is important for KP LUAD growth in vivo.

Due to the heterogeneity of HNF4α expression in the KP model in vivo, we generated organoids from our KPH GEMM and derived subclones (Pleguezuelos-Manzano et al., 2020) in which NKX2-1 and HNF4α were uniformly co-expressed, hereafter referred to as dual-positive (DP) organoids (**Figure 1F**). Despite evidence that the gastric-like NKX2-1-positive state is highly plastic (LaFave et al., 2020; Marjanovic et al., 2020; Yang et al., 2022), NKX2-1/HNF4α co-expression was stable, which enabled us to directly assess the cell-autonomous consequences of *Hnf4a* deletion. Treatment of these organoids in vitro with 4-hydroxytamoxifen (4OHT) generates isogenic organoid pairs that differ only in HNF4α expression (**Figure 1F, Figure S1C**). Importantly, *Hnf4a* deletion in DP organoids strikingly and significantly slowed overall growth (**Figure 1G**), which was accompanied by both significantly increased cell death (**Figure S1D**) and impaired proliferation (**Figure S1E**). We also treated DP organoids with BI6015, an inhibitor of HNF4α which binds to the ligand binding pocket and dampens HNF4α transcriptional activity and DNA binding (Kiselyuk et al., 2012). Pharmacologic inhibition of HNF4α with BI6015 decreased growth of DP organoids and attenuated the expression of HNF4α target genes in a dose-dependent manner (**Figure S1F-G**). Consistent with these data, we found that exogenous HNF4α enhances in vivo growth of the DP human LUAD cell line H1651 (**Figure S1H-I**) as assessed by tumor volume, mass and mitosis rate (**Figure 1H-K)**. In conclusion, analysis of primary human LUAD in conjunction with in vitro and in vivo models reveals that HNF4α can significantly enhance the growth of NKX2-1-positive LUAD.

### HNF4α induces a gastrointestinal/hepatic differentiation state in NKX2-1-positive LUAD

HNF4α is a robust marker of the previously defined gastric-like state in NKX2-1-positive LUAD (LaFave et al., 2020; Marjanovic et al., 2020; Yang et al., 2022) and regulates differentiation in the normal gastrointestinal (GI) tract and liver (Alder et al., 2014; Babeu & Boudreau, 2014; Chen et al., 2020; Garrison et al., 2006; Girard et al., 2022; Li et al., 2000). We therefore asked whether HNF4α directly activates GI and/or hepatic differentiation programs in NKX2-1-positive LUAD. To address this question, we first performed differential gene expression analysis on *HNF4A*-high vs *HNF4A*-low patients from our curated LUAD TCGA patient cohort (**Supplemental Tables S1 and S2**). GSEA pathway analysis revealed enrichment of GI and liver-like cell identity signatures in the *HNF4Α*-high vs *HNF4Α*-low group (**Figure 2A, Supplemental Table S3**). Additionally, *HNF4A* expression was positively correlated with several GI HNF4α target genes including *LGALS4*, *EPS8L3* and *HNF1Α* in the TCGA cohort (**Figure S2A**).

**Figure 2.**
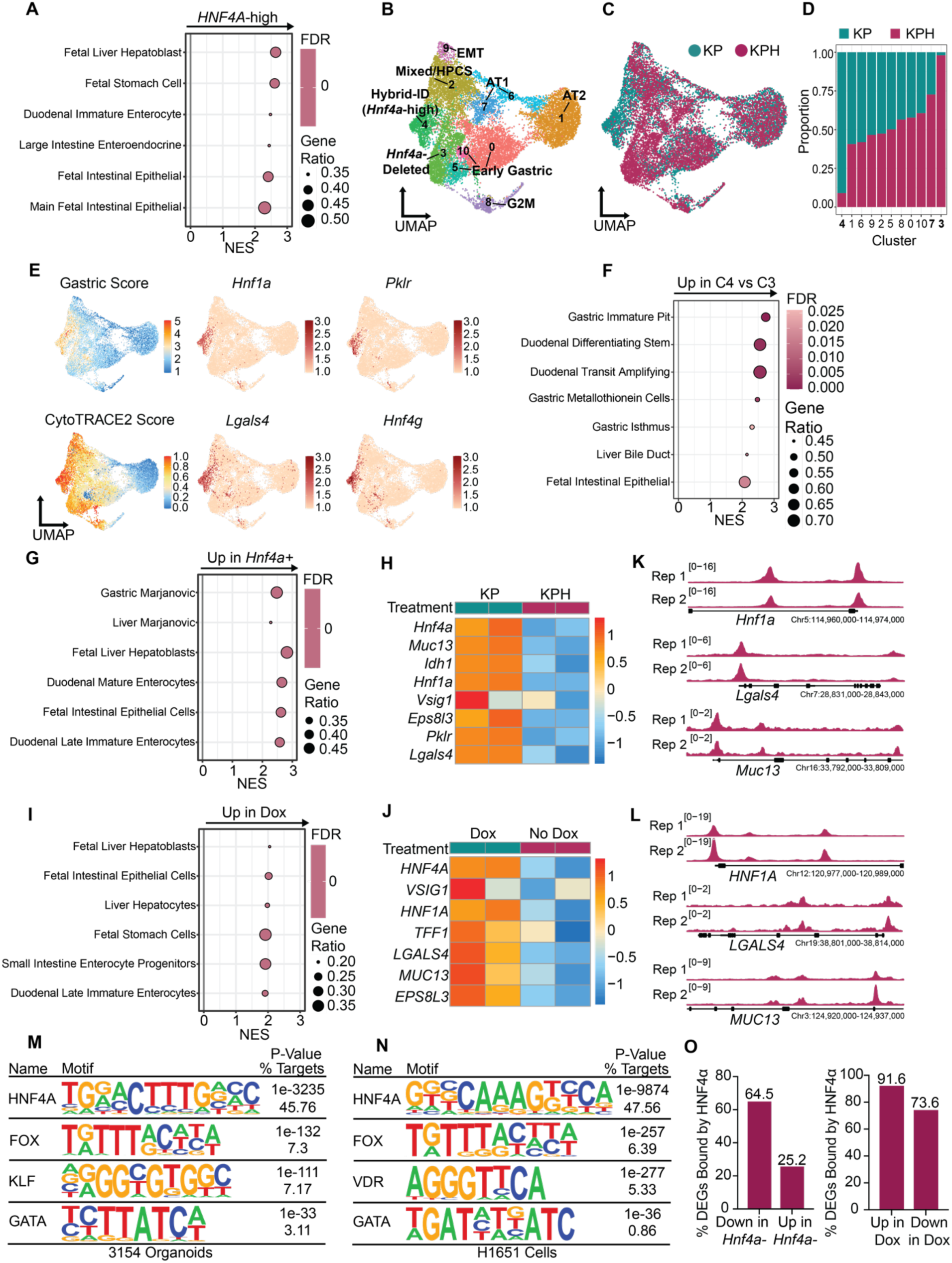
HNF4α drives gastrointestinal and liver-like differentiation programs within NKX2-1-positive LUAD. **A.** GSEA pathway analysis of differentially expressed genes in *HNF4A*-high vs *HNF4A*-low TCGA LUAD tumors showing representative cell type signatures. **B-F**. scRNA-seq was performed on tumor cells isolated from KP or KPH mice. Mice were treated with tamoxifen or vehicle control 10 weeks post-tumor initiation with adenoviral mSPC-FlpO and collected 2 weeks later. **B.** UMAP of KP (n=6823) and KPH (n=7544) tumor cells colored by Seurat-defined clusters. **C.** UMAP of KP and KPH tumor cells colored by tumor genotype. **D.** Proportion of tumor cells from each genotype per Seurat cluster. **E.** Relative expression of gastric gene signature, differentiation score from CytoTRACE2 analysis, and imputed expression of HNF4α target genes. **F.** GSEA pathway analysis on differentially expressed genes in cluster 3 (KPH-specific) vs cluster 4 (KP-specific) showing representative cell type signatures. **G.** GSEA pathway analysis on differentially expressed genes in control vs *Hnf4a*-deleted (2 weeks-post 4OHT treatment) 3154 DP organoid line showing representative cell type signatures. **H.** Heatmap showing expression of selected HNF4α target genes in control vs *Hnf4a*-deleted 3154 DP organoid line. Z-scores across samples of log2 normalized counts shown. **I.** GSEA pathway analysis on differentially expressed genes in H1651 cell line harboring doxycycline-inducible HNF4α overexpression construct showing representative cell type signatures. Cells were treated with 1μg/mL doxycycline to induce exogenous HNF4α (or vehicle control) for 7 days. **J.** Heatmap showing expression of selected HNF4α target genes in H1651 cell line treated with dox to overexpress HNF4α or control. Z-scores across samples of log2 normalized counts shown. **K.** ChIP-seq tracks illustrating HNF4α binding at gastrointestinal target genes in 3154 murine DP organoid line (n=2 biological replicates). **L.** ChIP-seq tracks illustrating HNF4α binding at gastrointestinal target genes in HNF4α-overexpressing H1651 cells (n=2 biological replicates). **M.** HOMER motif enrichment analysis of intersected HNF4α ChIP-seq peaks from two biological replicates in 3154 DP murine organoid line. **N.** HOMER motif enrichment analysis of intersected ΗNF4α ChIP-seq peaks from two biological replicates in H1651 HNF4α OE cell line. **O.** Percentage of significant differentially expressed genes (padj < 0.05, log2FC > 0.585 or < −0.585) in 3154 organoids or H1651 cells that overlap with genes bound by HNF4α.

We next asked which cell identity states were functionally dependent on expression of HNF4α in vivo. We deleted *Hnf4a* in our KPH GEMM at 10 weeks post tumor initiation, a timepoint at which a significant number of tumor cells are HNF4α-positive (**Figure 1C**) and performed scRNA-seq on tumor cells from KP (n=2) and KPH (n=2) mice 2 weeks later (12 weeks post initiation). Unsupervised clustering of single-cell transcriptomes revealed several distinct clusters representing stromal cells, immune cells, normal lung cells, and transformed LUAD cells (**Figure S2B-C, Supplemental Table S4**). We did not note any major differences in clustering between KP vs KPH cells within stromal and immune populations, indicating that our datasets did not contain major batch effects and that tumor-specific *Hnf4a* deletion had little effect on the transcriptomes of surrounding cell populations. Reclustering of only tumor cells identified by high expression of the *Cre^ERT2^* transgene and low expression of stromal-specific markers (**Figure S2C**) revealed 11 distinct clusters, of which clusters 3 and 7 were comprised of mostly KPH tumor cells and cluster 4 was predominated by KP control tumor cells (**Figure 2B-D, Supplemental Table S4**). Analysis of marker genes of each tumor cell cluster as well as application of cell-type signature scores derived from previous studies enabled us to correlate these clusters to previously defined cell identity programs in the KP mouse model (LaFave et al., 2020; Li et al., 2024; Marjanovic et al., 2020; Yang et al., 2022) (**Figure 2B, Figure S2F, Supplemental Table S4**). Importantly, quantitation of floxed (exons 4 and 5) vs non-floxed exons revealed successful recombination of *Hnf4a* in KPH tumor cells overall (**Figure S2D, Supplemental Table S4**). Additionally, this analysis enabled us to infer which cells likely expressed HNF4α prior to tamoxifen administration in our GEMM, and thus which cell states are dependent on HNF4α for maintenance. AT2-like cluster 1 and AT1-like cluster 6 had either undetectable or minimal expression of floxed and non-floxed *Hnf4a* exons in both KP and KPH cells, reflecting the early *Hnf4a*-negative status of these states during normal LUAD evolution (**Figure S2E**). In KPH-specific clusters 3 and 7, however, non-floxed exons of Hnf4a were expressed, but expression of floxed exons was completely ablated in these clusters, in contrast to surrounding KP-predominated clusters (**Figure S2E**), indicating that clusters 3 and 7 represent de novo differentiation states with a history of *Hnf4a* expression to which tumor cells may equilibrate following deletion.

Strikingly, expression of the gastric-like program and individual GI marker genes including *Hnf1a*, *Lgals4*, *Hnf4g*, and *Pklr* were largely restricted to KP-dominated cluster 4 and were significantly lower in neighboring KPH cluster 3 (**Figure 2E**). Additionally, CytoTRACE2, a computational tool that infers differentiation state based on the number of uniquely expressed genes per cell (Gulati et al., 2020), predicted that gastric-like cluster 4 was the most dedifferentiated cluster in our dataset. In contrast, KPH-specific cluster 3 was relatively more differentiated, indicating that *Hnf4a* deletion may induce a partial redifferentiation of KPH tumor cells (**Figure 2E**). Finally, pathway analysis of genes differentially expressed between KP-specific cluster 4 and KPH-specific cluster 3 revealed that cluster 4 was enriched for gene signatures associated with GI-like and liver-like identity programs (**Figure 2F**). Notably, cluster 4 was also enriched for gene signatures associated with cellular growth and proliferation, reflecting the pro-tumor effects of HNF4α that we observe in the KP model (**Supplemental Table S4**). We also turned our attention to the previously-described High Plasticity Cell State (HPCS) that has been implicated as a highly plastic, aggressive and therapy resistant state in KP tumor evolution (LaFave et al., 2020; Li et al., 2024; Marjanovic et al., 2020; Yang et al., 2022). Interestingly, we find high overall expression of *Hnf4a* in the HPCS cluster, but *Hnf4a*-deleted HPCS cells still cluster closely to *Hnf4a*-proficient KP HPCS cells, suggesting that HNF4α may not play an important functional role within the HPCS state (**Figure S2H**).

We also performed bulk RNA-seq on two in vitro models of DP LUAD; a KPH organoid line derived from our GEMM (3154 organoids) and a human LUAD cell line (H1651 cells). Differential gene expression analysis revealed 765 upregulated and 1179 downregulated genes upon *Hnf4a* deletion in KPH organoids (**Supplemental Table S2).** GSEA pathway enrichment analysis revealed an enrichment for GI and liver-related cell and tissue type gene signatures in KP vs KPH organoids (**Figure 2G, Supplemental Table S3**). Additionally, expression of several individual GI-like HNF4α target genes was attenuated following *Hnf4a* deletion in DP organoids (**Figure 2H**). In H1651 DP human cells, exogenous HNF4α induced 179 genes and repressed 79 genes (**Supplemental Table S2).** Like the murine organoid line, GSEA pathway analysis also revealed enrichment for GI and liver-like gene signatures in *HNF4A*-high vs *HNF4A*-low H1651 cells as well as increased expression of specific GI HNF4α target genes upon HNF4α overexpression (**Figure 2I-J, Supplemental Tables S2 and S3**). Finally, GSEA pathway analysis using a curated list of published HNF4α regulated genes from normal tissues or other cancer types revealed a strong positive enrichment of these signatures in HNF4α-high organoids and human cells (**Figure S2G**) (Camolotto et al., 2021; Chen et al., 2019; Garrison et al., 2006; Sumi et al., 2007; Thakur et al., 2024; Thymiakou et al., 2020; Xu et al., 2020). In both our murine and human models, the genes driving top enriched liver versus GI-like signature scores displayed less than 50% overlap, indicating that HNF4α drives multiple distinct cell identity programs at once within DP LUAD, ultimately leading to a hybrid-ID state. Of note, we also observed this pattern in our pathway enrichment results from the TCGA *HNF4A* cohorts. Collectively, our scRNA-seq and bulk RNA-seq experiments reveal that HNF4α activates canonical GI and liver-like differentiation programs when aberrantly expressed within the lung, a tissue normally devoid of HNF4α expression.

Next, we asked where HNF4α can bind throughout the genome when expressed in the foreign epigenetic context of the lung. We performed chromatin immunoprecipitation sequencing (ChIP-seq) for HNF4α on DP murine organoids as well as human cells treated with doxycycline to overexpress HNF4α. We identified 10,684 HNF4α ChIP-seq peaks in the DP organoids and 53,286 HNF4α ChIP-seq peaks in the DP human cell line and annotated these peaks to the nearest gene TSS (**Supplemental Table S5**). We found that HNF4α directly binds near many canonical GI target genes in both human and murine hybrid-ID models including *HNF1A*, *LGALS4*, and *MUC13* (**Figure 2K-L**). HOMER motif enrichment analysis revealed that the HNF4α motif was the top enriched motif in HNF4α-bound regions in these datasets, followed by other transcription factors known to interact or coordinate with HNF4α in different normal tissue and disease contexts, including FOX, KLF, and GATA family member proteins (**Figure 2M-N**) (Alder et al., 2014; Chahar et al., 2014; San Roman et al., 2015; Snyder et al., 2013; Wallerman et al., 2009). We also intersected HNF4α-bound regions with published HNF4α ChIP-seq datasets from normal murine liver (Qu et al., 2021), duodenum (Chen et al., 2019), and colon tissue (Gu et al., 2024), which revealed significant overlap between these datasets and our HNF4α ChIP-seq datasets (**Figure S2I**). To further characterize HNF4α activity within hybrid-ID LUAD, we performed an integrative analysis of our bulk RNA-seq and ChIP-seq datasets. In the DP organoids, HNF4α binds near the majority of genes significantly downregulated following *Hnf4a* deletion, but a smaller proportion of genes induced upon *Hnf4a* deletion were bound by HNF4α (**Figure 2O**). Likewise, in H1651 cells, a larger proportion of genes significantly induced upon HNF4α ΟΕ were bound by HNF4α than genes that were downregulated following HNF4α OE (**Figure 2O**). Together, these integrated analyses suggest that HNF4α functions primarily as a transcriptional activator in hybrid-ID LUAD. Finally, we derived a consensus set of HNF4α-regulated genes by combining shared differentially expressed genes positively regulated by HNF4α in both human and murine bulk RNA-seq datasets (**Supplemental Table S3**). We applied a gene score for this HNF4α-regulated signature to each sample in our curated TCGA cohort using single-sample GSEA (ssGSEA). We found that enrichment of the consensus HNF4α-regulated gene signature was positively correlated with expression of *HNF4A* in this dataset (**Figure S2J**). Together, these results show that HNF4α directly binds and activates genes associated with distinct non-pulmonary differentiation states, including GI and liver-like programs, to induce a hybrid identity state in NKX2-1-positive LUAD.

Cellular identity is defined not only by RNA expression patterns but also by functional differences between cells. Further analysis of our bulk RNA-seq datasets revealed that, in addition to activating GI/liver identity programs, HNF4α also induced multiple metabolism-associated gene expression pathways (**Figure S3A-B, Supplemental Table S3**). Individual genes associated with various arms of metabolism including glucose metabolism and lipid metabolism were downregulated upon *Hnf4a* deletion in DP LUAD organoids (**Figure S3C**). Since nutrient utilization and metabolism also display tissue-specific patterns, we asked whether HNF4α modulates metabolism in hybrid-ID LUAD. To perform an unbiased analysis of changes in metabolite levels in DP organoids following *Hnf4a* deletion, we performed gas chromatography-mass spectrometry (GC-MS)-based metabolomics and liquid chromatography-MS (LC-MS)-based lipidomics on isogenic KP and KPH murine organoids. *Hnf4a* deletion caused widespread disruption of the metabolome, with alterations to metabolites related to central carbon metabolism, fructose metabolism, and amino acid metabolism, among other pathways (**Figure S3D-E, Supplemental Table S7**). We also observed a striking lipid accumulation phenotype following *Hnf4a* deletion in our lipidomics dataset (**Figure S3F-G, Supplemental Table S7**). This may reflect the canonical role of HNF4α in regulating lipid metabolism in normal tissues, including the intestines and the liver (Chen et al., 2020; Huang et al., 2020).

### HNF4α suppresses pulmonary differentiation programs in NKX2-1-positive LUAD

We next sought to characterize the identity adopted by HNF4α-low cells in our LUAD models. GSEA pathway analysis of enriched programs in *HNF4A*-low vs *HNF4A*-high TCGA LUAD samples revealed enrichment for pulmonary cell type signatures, including AT1 and AT2-like gene signatures (**Figure 3A, Supplemental Table S3**). Since HNF4α is never expressed in the lung or in precursor cells of the lung during development (Duncan et al., 1994; Islam et al., 2024; Vincent & Robertson, 2004), our finding that HNF4α may influence pulmonary differentiation within hybrid-ID LUAD was unexpected. Nevertheless, we also observed changes to alveolar/pulmonary programs and marker genes in our functional studies. In our scRNA-seq dataset, KPH-specific cluster 7 was highly enriched for AT1-like markers including *Hopx* and *Ager* and a gene signature of AT1 cells (Du et al., 2015) (**Figure 3B**). Moreover, KPH-specific cluster 3 highly expressed AT2-like markers including *Cxcl15* and *Sftpc* as well as an AT2-specific gene signature (Marjanovic et al., 2020) (**Figure 3B**). GSEA pathway analysis of differentially expressed genes between cluster 3 and cluster 4 revealed an enrichment of alveolar and pulmonary gene signatures in KPH-specific cluster 3 compared to the hybrid-ID, KP-specific cluster 4 (**Figure 3C, Supplemental Table S4**). Of note, AT2-like cluster 1 expressed higher levels of canonical AT2 marker genes including *Lyz2* and *Sftpa1* than cluster 3, suggesting that deletion of *Hnf4a* does not completely revert cells back to an AT2-high state adopted by early low-grade cells in the KP model (**Supplemental Table S4**). Additionally, comparison of cluster 6 (KP/KPH AT1-like) and cluster 7 (KPH-specific AT1-like) revealed that cluster 6 more highly expresses AT2 marker genes, suggesting that deletion of *Hnf4a* may induce a more exclusive AT1 identity in these cells rather than more mixed AT1/AT2 differentiation states that naturally emerge during KP tumor evolution (**Supplemental Table S4**). *Hnf4a* deletion also induces alveolar differentiation programs and marker genes in murine DP organoids (**Figure 3D-E, Supplemental Table S3**). Moreover, exogenous HNF4α expression in two DP organoid lines significantly reduced expression of the canonical AT2 and AT1 cell markers *Sftpc* and *Ager* (**Figure 3F-G**). Overexpression of HNF4α in H1651 cells also attenuated expression of NKX2-1 targets and alveolar marker genes including *LMO3* and *SPOCK1* (**Figure 3H**). Together, these results support a non-canonical role of HNF4α in the suppression of pulmonary identity programs when it is co-expressed with NKX2-1 in the context of hybrid-ID LUAD.

**Figure 3.**
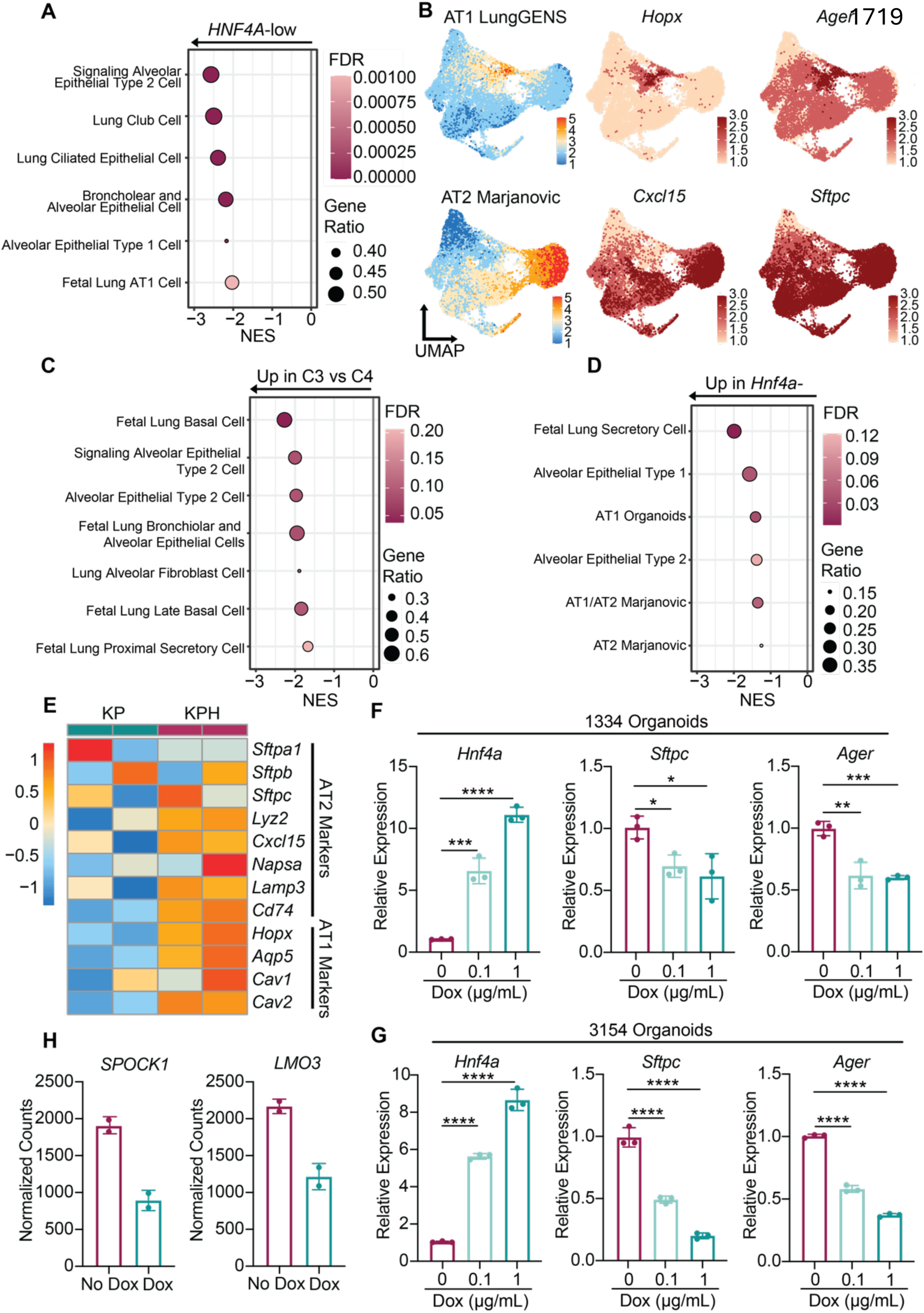
HNF4α suppresses alveolar differentiation programs in NKX2-1-positive LUAD. **A.** GSEA pathway analysis of differentially expressed genes in *HNF4A*-high vs *HNF4A*-low TCGA patients showing representative cell type signatures. **B.** UMAPs showing relative expression of alveolar gene signatures and specific markers of AT1 and AT2 cells. **C.** GSEA pathway analysis on differentially expressed genes in cluster 3 (KPH-specific) vs cluster 4 (KP-specific) showing representative cell type signatures. **D.** GSEA pathway analysis on differentially expressed genes in 3154 control vs *Hnf4a*-deleted DP organoid line showing representative cell type signatures. **E.** Heatmap showing expression of selected pulmonary marker genes in 3154 control vs *Hnf4a*-deleted DP organoid line. Z-scores across samples of log2 normalized counts shown. **F-G.** 1334 (F) and 3154 (G) DP organoid lines were transduced with HNF4α overexpression constructs and treated with two concentrations of doxycycline. RT-qPCR illustrating expression of *Hnf4a*, *Sftpc* (AT2 marker) and *Ager* (AT1 marker) following overexpression of HNF4α (one representative replicate shown of n=2 biological replicates; one-way ANOVA with Dunnett’s multiple comparisons test, *p<0.05, **p<0.002, ***p<0.001, ****p<0.0001; error bars, SD). **H.** Normalized expression of NKX2-1 target genes *SPOCK1* and *LMO3* in dox-inducible HNF4α-overexpression cell line (H1651) by bulk RNA-seq.

### HNF4α interacts with NKX2-1 in hybrid-ID LUAD

We next sought to determine the molecular mechanisms by which HNF4α disrupts pulmonary differentiation states in hybrid-ID LUAD. We hypothesized that HNF4α may dampen expression of pulmonary identity programs by disrupting activity and/or genomic binding of key pulmonary lineage specifiers. We turned our attention to the master lung lineage specifier NKX2-1 and performed ChIP-seq for NKX2-1 as well as the pioneer factors FoxA1 and FoxA2. Importantly, FoxA1/2 bind with NKX2-1 during normal lung differentiation as well as in LUAD (Little et al., 2021; Minoo et al., 2007; Snyder et al., 2013; Watanabe et al., 2013). Moreover, FoxA1/2 can also both activate and bind with HNF4α in order to induce hepatic and GI-like differentiation in normal tissue and in LUAD (Alder et al., 2014; Camolotto et al., 2018; Iwafuchi-Doi et al., 2016; Orstad et al., 2022). In line with the normally dichotomous roles of NKX2-1 and HNF4α, analysis of publicly available HNF4α ChIP-seq data from the murine liver (Qu et al., 2021), duodenum (Chen et al., 2019), and colon (Gu et al., 2024) revealed that in these tissues, HNF4α strongly binds to the GI HNF4α target gene *Hnf1a* but not to pulmonary target genes including *Sftpb* (**Figure S4A**). Likewise, analysis of NKX2-1 binding patterns in normal murine AT1 and AT2 cells (Little et al., 2021) revealed binding of NKX2-1 to *Sftpb* but not to *Hnf1a* (**Figure S4A**). Additionally, comparison of HNF4α binding sites in the duodenum, liver, or colon vs NKX2-1 binding sites in AT1 or AT2 cells revealed minimal overlap (**Figure S4B**). Further, in a p53-proficient model of LUAD, NKX2-1 binds to pulmonary genes including *Sftpb* but not to GI target genes (Snyder et al., 2013). Surprisingly, in stark contrast to these normal binding patterns, we observed extensive overlap in genome-wide binding of NKX2-1, HNF4α, and FoxA1/2 in DP human cells and murine organoids (**Figure 4A-B, Figure S4C-D, Supplemental Table S5**). This overlap in binding was not due to a lack of specificity, as motif enrichment analysis revealed NKX2 and FOX motifs among the top most enriched motifs in our NKX2-1 and FoxA1/2 ChIP-seq experiments, respectively (**Supplemental Table S5**). NKX2-1 and HNF4α, as well as FoxA1/2, bind to both canonical NKX2-1 pulmonary targets such as *Sftpb* and *S*ftpa1 as well as GI HNF4α target genes including *Lgals4* and *Hnf1a* (**Figure 4C-D**). Together, these observations support the notion that the observed colocalization between NKX2-1 and HNF4α reflects aberrant patterns of transcription factor binding that diverges from normal biology and represents a unique characteristic of the hybrid-ID state in LUAD.

**Figure 4.**
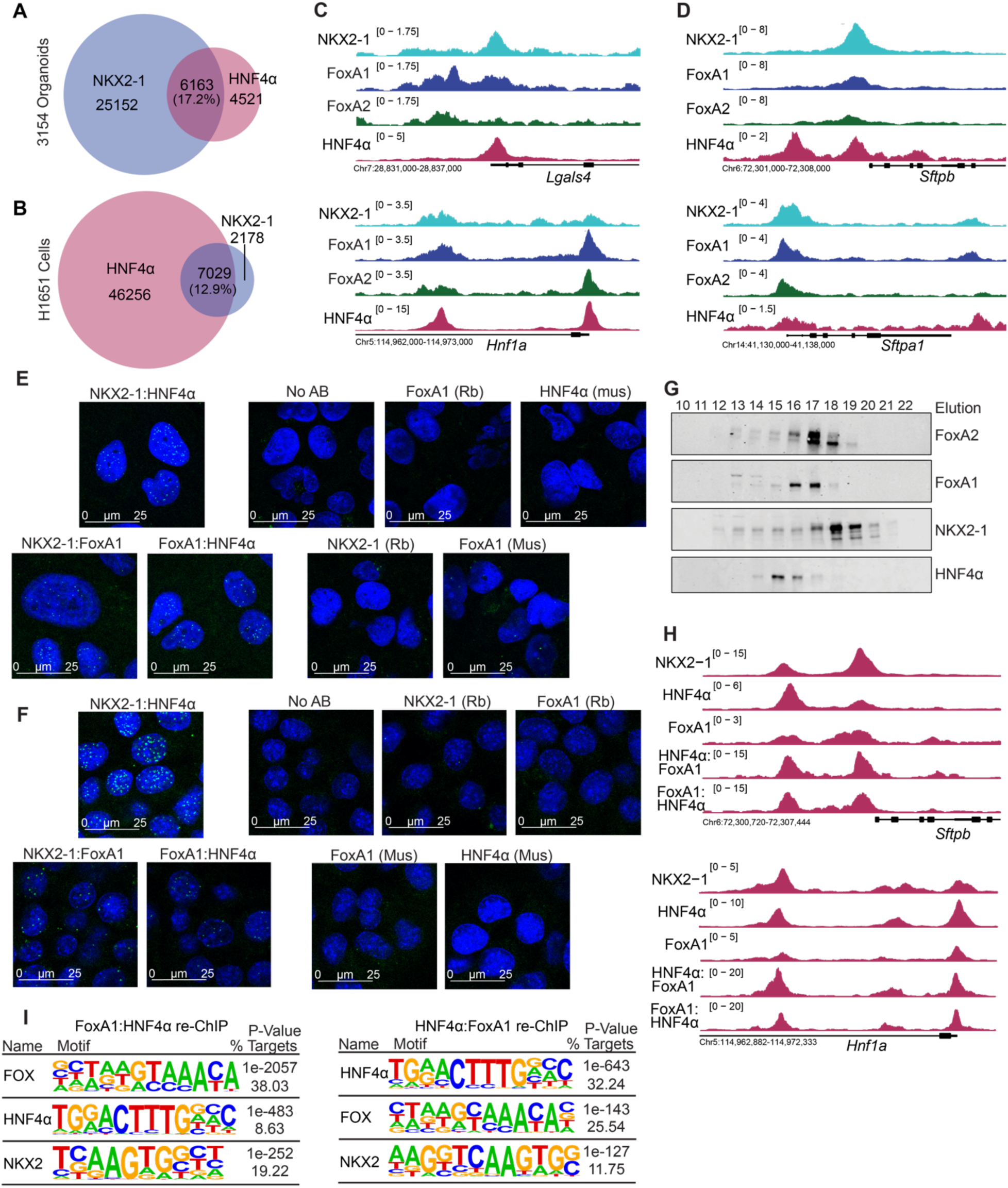
HNF4α associates with NKX2-1 and FoxA1/2 in hybrid-identity LUAD. **A.** Quantification of overlapping peaks (number of overlapping peaks and percent overlap out of total peaks) between NKX2-1 and HNF4α ChIP-seq datasets in 3154 organoids **B.** Quantification of overlapping peaks (number of overlapping peaks and percent overlap out of total peaks) between NKX2-1 and HNF4α ChIP-seq datasets in H1651 cells. **C.** ChIP-seq tracks showing binding of NKX2-1, FoxA1, FoxA2, and HNF4α near the TSS of gastrointestinal genes *Lgals4* and *Hnf1a* in 3154 organoids. **D.** ChIP-seq tracks showing binding of NKX2-1, FoxA1, FoxA2, and HNF4α near the TSS of pulmonary genes *Sftpb* and *Sftpa1* in 3154 organoids. **E.** PLA assay in fixed H1651 human cells stained with DAPI (blue) showing no-antibody and single antibody only controls, and nuclear PLA signal (green) for NKX2-1:FoxA1, HNF4α:FoxA1, and NKX2-1:HNF4α. (Scale bar = 25μM. Representative images shown from n=2 biological replicates). **F.** PLA assay in 3311 DP cell line stained with DAPI (blue) showing no-antibody and single antibody controls and nuclear PLA signal (green) for NKX2-1:FoxA1, HNF4α:FoxA1, and NKX2-1:HNF4α (Scale bar = 25μM. Representative images shown from n=2 biological replicates). **G.** Gel filtration chromatography experiment in 3311 DP murine cell line (one representative replicate shown of n=2 biological replicates). **H.** ChIP-seq tracks of FoxA1, HNF4α, and NKX2-1 as well as FoxA1:HNF4α and HNF4α:FoxA1 sequential ChIP experiments in 3311 DP murine cells showing binding of FoxA1 and HNF4α at *Hnf1a* and *Sftpb* (one representative replicate shown of n=2 biological replicates). **I.** HOMER motif enrichment analysis for FoxA1:HNF4α and HNF4α:FoxA1 sequential ChIP experiments showing enrichment for FOX, HNF, and NKX motifs at FoxA1:HNF4α co-bound sites.

Overlapping binding sites between HNF4α and NKX2-1 might reflect formation of an aberrant protein complex that diverges from normally dichotomous roles and binding patterns of these transcription factors in differentiation and tissue specification. Alternatively, overlapping binding sites could reflect mutually exclusive, and potentially antagonistic, binding of HNF4α and NKX2-1 in distinct subpopulations of cells. We therefore performed proximity ligation assays (PLA) on DP human and murine cell lines to assess physical interactions between NKX2-1, HNF4α, and FoxA1 in individual cells. PLA experiments revealed expected nuclear-localized FoxA1-HNF4α and FoxA1-NKX2-1 interactions in both human and murine cells (**Figure 4E-F**). Importantly, we also observed nuclear-localized interactions between NKX2-1 and HNF4α in both cell lines, indicating that NKX2-1, HNF4α and FoxA1 form a previously uncharacterized protein complex within hybrid-ID LUAD (**Figure 4E-F**).

As a complementary approach, we performed gel filtration chromatography on nuclear-enriched lysates from KP DP murine cells. This technique enables the separation of proteins and protein complexes by both size and shape. Notably, we observed that NKX2-1, HNF4α, FoxA1 and FoxA2 elute in fractions corresponding to molecular weights larger than their monomeric or, where applicable, dimeric forms, suggesting that they likely exist in complexes with additional proteins (**Figure 4G**). Consistent with our PLA results, NKX2-1 co-elutes with both HNF4α and FoxA1/2 in several fractions. Previous work from our group has revealed that NKX2-1 binds with FoxA1/2 to activate a large proportion of NKX2-1 target genes in LUAD, but that NKX2-1 can also bind and activate a subset of target genes in a FoxA1/2-independent manner (Orstad et al., 2022). The presence of FoxA1/2-negative NKX2-1 fractions thus support the notion of that a pool of NKX2-1 exists in alternate protein complexes which do not contain FoxA1/2 (**Figure 4G**). Additionally, the presence of NKX2-1-containing fractions with no HNF4α protein suggest that only a subset of NKX2-1 molecules present within hybrid-ID LUAD complex with HNF4α (**Figure 4G**).

Finally, we optimized a sequential ChIP-seq (ChIP-re-ChIP) protocol (Seneviratne et al., 2024) to assess HNF4α and FoxA1 co-occupancy at pulmonary and GI-specific sites in NKX2-1-positive LUAD. Analysis of overlapping peaks called in two biological replicates of FoxA1:HNF4α ChIP-re-ChIP revealed 13,787 co-bound peaks, while HNF4α:FoxA1 ChIP-re-ChIP experiments resolved 2,871 peaks (**Supplemental Table S8**). Consistent with their binding patterns in normal tissues, both datasets showed enrichment at GI targets including *Hnf1a* and *Lgals4* (**Figure 4H**). However, we also observed co-binding of HNF4α and FoxA1 at pulmonary genes, including *Sftpb*, indicating that HNF4α and FoxA1 bind together at both GI and pulmonary marker genes in hybrid-ID LUAD (**Figure 4H**). Intersection of FoxA1 and HNF4α individual ChIP-seq peaks and ChIP-re-ChIP peaks revealed substantial overlap, reflecting the extensive colocalization of these two transcription factors in hybrid-ID LUAD (**Figure S4E**). Moreover, HOMER motif enrichment analysis revealed enrichment for HNF4, FOXA, and NKX motifs in HNF4α:FoxA1 co-bound regions, consistent with the possibility of a chromatin-interacting NKX2-1:HNF4α:FoxA1/2 protein complex within NKX2-1-positive LUAD (**Figure 4I**). Together, these experiments uncover a novel interaction between NKX2-1 and HNF4α in hybrid-ID LUAD that may contribute to the specification of cell identity programs in this LUAD subtype.

### HNF4α modulates cell identity-specific binding of NKX2-1

Based on our discovery of a protein complex containing both NKX2-1 and HNF4α in hybrid-ID LUAD, we hypothesized that HNF4α may physically sequester a subset of NKX2-1 molecules to GI genes, leaving a smaller pool of NKX2-1 to bind and activate canonical alveolar target genes, thus dampening pulmonary differentiation programs. To test this hypothesis, we performed ChIP-seq for NKX2-1 in *Hnf4a*-deleted organoids and compared binding patterns of NKX2-1 to *Hnf4a*-positive, control organoids. Differential binding analysis (DiffBind) revealed 476 gained and 316 lost NKX2-1 peaks following *Hnf4a* deletion, showing that HNF4α significantly alters NKX2-1 chromatin binding in hybrid-ID LUAD, despite overall expression levels of *Nkx2-1* remaining constant (**Figure 5A, Supplemental Tables S2, S6**). Importantly, these de novo NKX2-1 binding events are mostly associated with gene activation, as integration of annotated differential binding sites with our bulk RNA-seq dataset revealed that a larger proportion of de novo NKX2-1-bound genes are upregulated following *Hnf4a* deletion than downregulated (**Figure S5A**). HOMER motif enrichment analysis of differentially bound NKX2-1 sites following *Hnf4a* deletion revealed enrichment for the HNF4 motif within NKX2-1 binding sites specific to *Hnf4a* positive organoids (**Figure 5B**). Further, Enrichr pathway analysis of differential NKX2-1-bound genes revealed enrichment for GI cell type signatures among genes bound by NKX2-1 in *Hnf4a*-positive organoids (**Figure 5C**). In agreement with this, binding of NKX2-1 to GI HNF4α target genes, such as *Lgals4* and *Hnf1a* was attenuated following *Hnf4a* deletion in DP murine organoids (**Figure 5D**). Analysis of gained NKX2-1 binding sites following *Hnf4a* deletion revealed that de novo NKX2-1 bound regions are enriched for FOX motifs and TEAD motifs, both known NKX2-1 binding partners (**Figure 5B**). Additionally, pulmonary cell type signatures were enriched within NKX2-1-bound genes specific to *Hnf4a*-deleted organoids (**Figure 5C**), and NKX2-1 shows increased binding to both its AT1 and AT2-specific targets (Little et al., 2021) following *Hnf4a* deletion (**Figure 5E**). NKX2-1 binding at HNF4α binding sites in the normal duodenum decreased upon *Hnf4a* deletion (**Figure 5E**) (Chen et al., 2019). Dynamic NKX2-1 binding sites also correlate with known changes in chromatin accessibility during KP LUAD progression (LaFave et al., 2020). Integration of single-cell ATAC-sequencing (scATAC-seq) data from the KP model with our differential NKX2-1 ChIP-seq peaks revealed greater chromatin accessibility at KPH-specific NKX2-1 binding sites within AT1 and AT2-like LUAD cells, but a reduced accessibility at KPH-specific NKX2-1 binding sites within gastric-like KP LUAD cells (**Figure S5B**).

**Figure 5.**
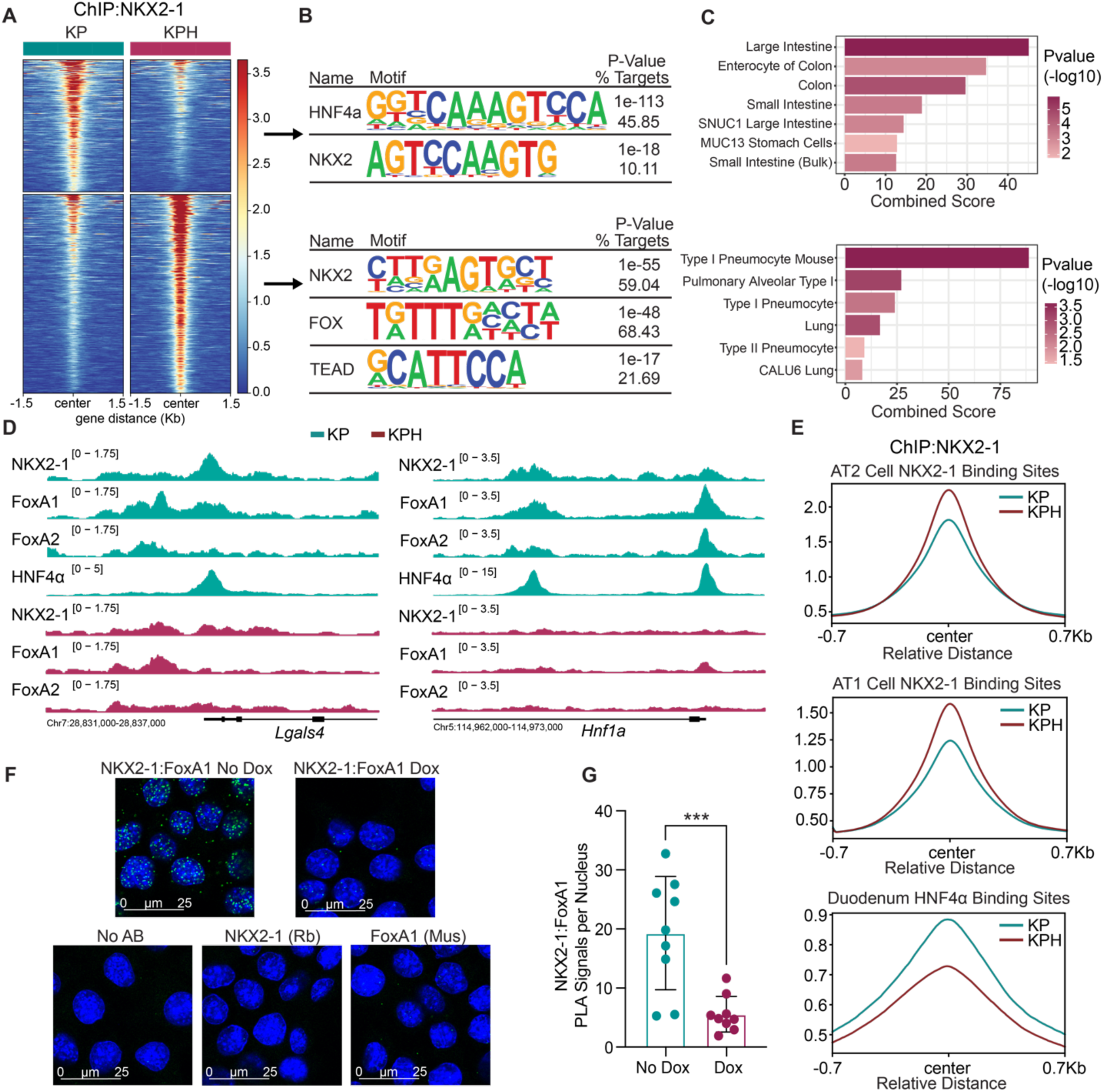
HNF4α modules cell identity-specific binding of NKX2-1. **A.** Heatmap depicting NKX2-1 occupancy at differential NKX2-1 binding sites following *Hnf4a* deletion in 3154 DP KPH organoid line. Differential binding analysis was performed using DiffBind with an adjusted p value cutoff of <0.05 and identified 316 KP-enriched peaks (out of 31,315 total KP NKX2-1 peaks) and 476 KPH-enriched peaks (out of 38,900 total KPH NKX2-1 peaks). **B.** HOMER motif enrichment analysis of differential NKX2-1 binding sites. **C.** Representative results from Enrichr pathway analysis of differential NKX2-1 binding sites annotated to closest TSS. **D.** ChIP-seq tracks showing co-binding of NKX2-1, FoxA1, FoxA2 and HNF4α to HNF4α target genes *Lgals4* and *Hnf1a* in control (green) and *Hnf4a*-deleted (red) 3154 organoids. **E.** Mean peak profile of control and *Hnf4a*-deleted NKX2-1 ChIP-seq signal at AT1-specific and AT2-specific NKX2-1 binding sites and duodenal HNF4α binding sites. **F.** PLA assay in 3311 DP cell line treated with 2.5μg/mL doxycycline to overexpress HNF4α (or vehicle control) stained with DAPI (blue) showing no-antibody and single antibody controls and nuclear PLA signal (green) for NKX2-1:FoxA1 (Scale bar = 25μM. Representative images shown from n=2 biological replicates). **G.** Quantification of NKX2-1:FoxA1 PLA signal from control and HNF4α OE 3311 cells, reported as average PLA signals per nucleus (quantification of n=4-5 fields of view from each of n=2 biological replicates; unpaired t-test p = 0.0008; error bars, SD).

In addition to NKX2-1, we found that *Hnf4a* deletion caused a similar change in FoxA1/2 binding. Specifically, FoxA1/2 relocalize towards pulmonary NKX2-1 target genes and away from GI-like HNF4α targets following *Hnf4a* deletion in DP organoids, even though overall expression of *Foxa1/2* does not change (**Figure 5D**, **Figure S5C-D, Supplemental Table S2**). Moreover, we performed a PLA experiment with a murine hybrid-ID cell line harboring a doxycycline-inducible HNF4α overexpression construct, and we found that overexpression of HNF4α significantly attenuated binding between NKX2-1 and FoxA1 (**Figure 5F-G, Figure S5E**).

In conclusion, these experiments reveal that HNF4α sequesters NKX2-1 away from canonical binding sites, which may reduce NKX2-1 binding with important cofactors involved in alveolar identity programs, including TEAD and FoxA1/2. These findings may provide a mechanistic explanation for the observed enrichment of AT1 and AT2 differentiation programs following *Hnf4a* deletion because NKX2-1 associates with FOX family members (specifically FoxA1 and FoxA2) during lung development and in LUAD to coordinately activate pulmonary identity (Minoo et al., 2007; Orstad et al., 2022; Snyder et al., 2013; Watanabe et al., 2013). NKX2-1 also associates with TEAD to induce AT1 differentiation programs within AT1 cells of the normal lung (Little et al., 2019; Little et al., 2021). Additionally, since NKX2-1 restrains growth in the KP model (Orstad et al., 2022; Snyder et al., 2013; Winslow et al., 2011), partial suppression of NKX2-1 activity is likely one mechanism by which HNF4α promotes tumor growth in NKX2-1-positive LUAD.

### RAS/MEK signaling maintains a hybrid-ID state by controlling NKX2-1 chromatin binding and activating *Hnf4a* expression

We have previously shown that FoxA1/2 activate HNF4α and other GI genes in KP LUAD to maintain a hybrid-ID state (Orstad et al., 2022). However, it is unclear why FoxA1/2 can activate a GI program in the presence of NKX2-1 in the KP model, but not the K model or normal lung epithelium (Snyder et al., 2013). We decided to investigate RAS/MEK signaling for multiple reasons. First, RAS/MEK signaling is associated with more aggressive and poorly differentiated disease in LUAD, and increased ERK activity can stimulate tumor progression (Cicchini et al., 2017; Feldser et al., 2010; Noguchi, 2010; Vicent et al., 2004). Specifically, grade 3 KP tumors demonstrate increased pERK as well as elevated HNF4α expression (Feldser et al., 2010). Recent studies suggest that MEK/ERK signaling may negatively regulate an alveolar identity in LUAD (Dost et al., 2020; Li et al., 2024; Maynard et al., 2020; Silberschmidt et al., 2011). In particular, RAS inhibitor treatment in the KP mouse model induces alveolar identity and reduces expression of ‘endodermal’ GI-like identity programs, but the mechanism underlying these transcriptomic changes is unknown (Li et al., 2024). Furthermore, these experiments were performed in a heterogeneous in vivo model, rendering it difficult to determine whether these concomitant identity changes occur in the same cells or reflect differential survival of distinct subpopulations following RAS inhibition (Li et al., 2024).

We hypothesized that the RAS/MEK signaling pathway either upstream of or in coordination with HNF4α may regulate hybrid-ID differentiation programs in NKX2-1-positive LUAD. To test this hypothesis, we performed RNA-seq on a murine DP cell line and organoid line treated with a MEK inhibitor (MEKi – Trametinib) and/or the KRAS^G12D^ inhibitor MRTX1133 (Wang et al., 2022) (**Supplemental Table S9**). Inhibition of MEK or KRAS^G12D^ induced AT1 and AT2-like cell type signatures and inhibited GI-like differentiation programs (**Figure 6A, Figure S6A, Supplemental Table S10**). Specifically, expression of several GI genes, including *Hnf4a* itself and many of its target genes (i.e. *Lgals4, Eps8l3, Vsig1*), were downregulated following RAS/MEK pathway inhibition, and specific markers of AT1 and AT2 identity (i.e. *Sftpc, Cd74, Ager, Pdpn*) were induced (**Figure 6B-C, Figure S6B**). We also found that our curated HNF4α-regulated gene signature was highly enriched in control organoids compared to MEKi or KRAS^G12D^ inhibitor treated cells (**Figure S6C**). Bulk-RNA seq on a human NKX2-1-positive/HNF4α-negative human cell line (H358) and a DP human cell line (H1651) also revealed an upregulation of AT1/AT2 marker genes and gene signatures following MEKi treatment (**Figure S6D-E, Supplemental Table S9**), suggesting that MEK activity can also dampen AT1/2 identity independent of HNF4α.

**Figure 6.**
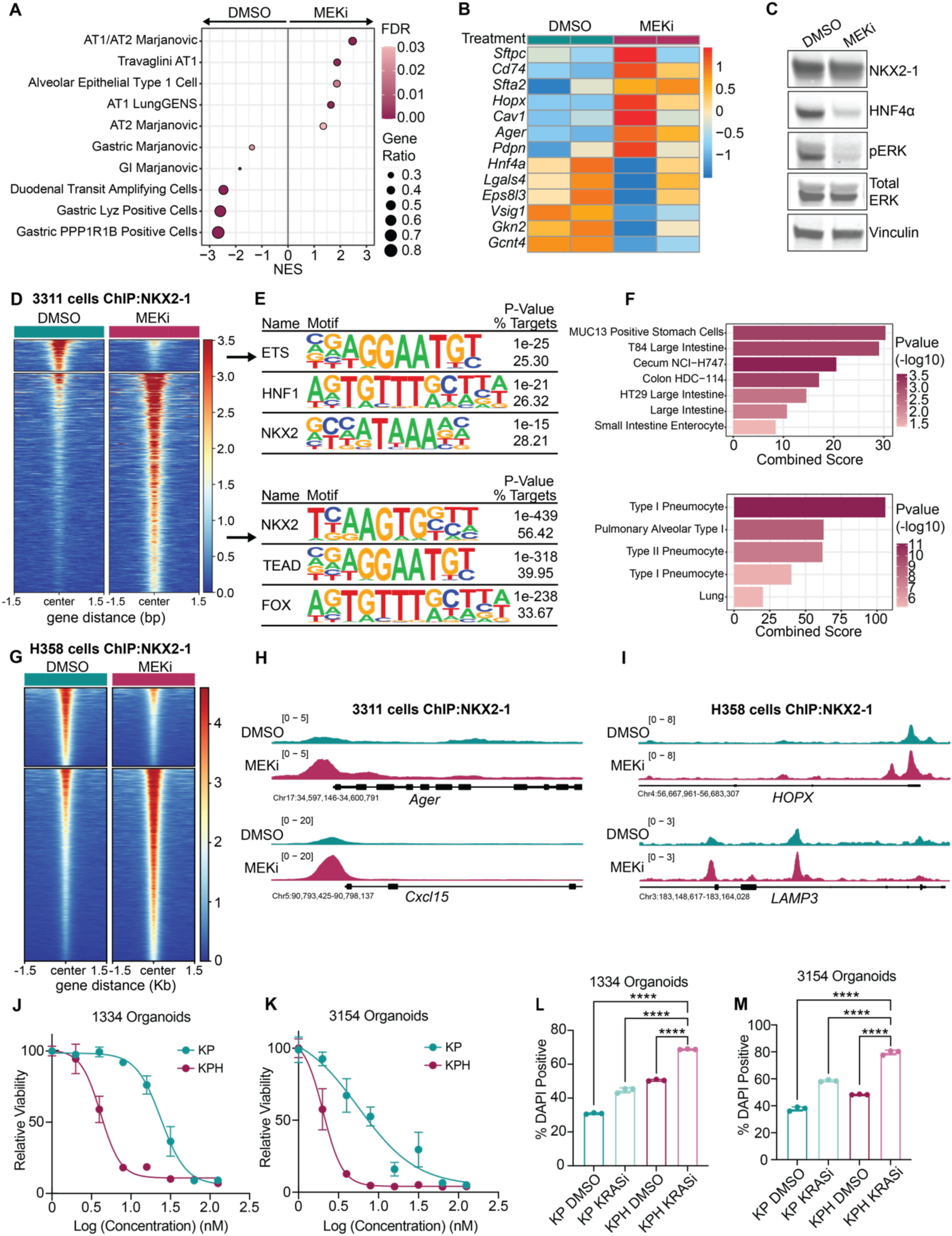
RAS/MEK signaling maintains a hybrid-ID state and controls NKX2-1 chromatin binding in LUAD. **A.** GSEA pathway analysis of differentially expressed genes in control vs MEKi-treated (50nM Trametinib) DP cell line (3311). Representative cell type gene signatures shown. **B.** Heatmap showing expression of selected gastrointestinal and pulmonary marker genes in control and MEKi-treated 3311 cells. Z-scores across samples of log2 normalized counts shown. **C.** Representative immunoblot of 3311 cells treated with 500nM Cobimetinib (MEKi). **D.** Heatmap depicting NKX2-1 occupancy at differential NKX2-1 binding sites following MEKi in DP cell line (3311). Differential binding analysis was performed using DiffBind with an adjusted p value cutoff of <0.05 which identified 706 control-specific peaks (out of 16,621 total control NKX2-1 peaks) and 4914 MEKi-specific peaks (out of 36,988 total MEKi NKX2-1 peaks). **E.** HOMER motif enrichment analysis of differential NKX2-1 binding sites. **F.** Enrichr pathway analysis of differential NKX2-1 binding sites annotated to closest TSS. **G.** Heatmap depicting NKX2-1 occupancy at differential NKX2-1 binding sites following MEKi in human H358 NKX2-1-positive cell line. Differential binding analysis was performed using DiffBind with an adjusted p value cutoff of <0.05 which identified 3041 control-specific peaks (out of 30,120 total control NKX2-1 peaks) and 7468 MEKi-specific peaks (out of 35,682 total MEKi NKX2-1 peaks). **H.** ChIP-seq tracks showing binding of NKX2-1 to AT1/2 targets in 3311 DP murine cells. **I.** ChIP-seq tracks showing binding of NKX2-1 to AT1/2 targets in H358 cells. **J-K.** Dose response curves of KP vs KPH DP organoid lines treated with MRTX1133 to inhibit KRAS^G12D^. (**J**) KP IC_50_=23.94nM, KPH IC_50_=4.138nM. (**K**) KP IC_50_=5.592nM, KPH IC_50_=2.034nM. (one representative replicate of n=2 biological replicates shown). **L-M**. DAPI staining for cell death in DP organoids treated with 3nM MRTX1133 for 72h (or vehicle control) and/or 4OHT to delete *Hnf4a*, measured by flow cytometry (one representative replicate shown of n=2 biological replicates; unpaired t test ****p<0.0001; error bars, SD).

Since RAS/MEK pathway inhibition in hybrid-ID LUAD partially phenocopies the effects of *Hnf4a* deletion (inhibition of GI programs and activation of alveolar identity), we hypothesized that this may also be due to changes in NKX2-1 chromatin binding. To test this, we profiled NKX2-1 binding in control and MEKi treated DP murine cells and organoids and resolved many differential NKX2-1 binding sites (706 control-specific peaks and 4914 MEKi-specific peaks in the hybrid-ID cell line and 7577 control-specific peaks and 3639 MEKi-specific peaks in the organoid line) (**Figure 6D, Figure S6F, Supplemental Table S6**). NKX2-1-bound sites specific to control cells were enriched for the NKX2 motif as well as HNF motifs (**Figure 6E).** ETS motifs, key downstream effectors of ERK signaling, were also enriched within NKX2-1-bound sites (**Figure 6E**). In MEKi-treated cells, NKX2-1 relocalized to sites enriched for TEAD and FOX motifs, potentially reflecting increased binding with these cofactors critical for induction of AT1 and AT2 identity programs (**Figure 6E**). Pathway analysis of differentially bound genes revealed enrichment of GI-like programs within control NKX2-1 binding sites and enrichment of AT1 and AT2 cell identity programs within de novo NKX2-1-bound genes following MEKi (**Figure 6F**). Importantly, we observed minimal difference in total levels of NKX2-1 itself following MEK inhibition in the DP murine cells, reinforcing the idea that a shift in NKX2-1 activity and/or binding, rather than levels, may be driving acquisition of alveolar programs (**Figure 6C**). We also profiled NKX2-1 binding in the NKX2-1-positive/HNF4α-negative H358 LUAD cell line +/- MEKi treatment, which revealed extensive relocalization of NKX2-1 following MEK inhibition (3041 control-specific peaks vs 7468 MEKi-specific peaks) (**Figure 6G, Supplemental Table S6**). Binding of NKX2-1 to AT1 and AT2-specific marker genes increased following MEK inhibition in all models tested (**Figure 6H-I, Figure S6G**). Further, NKX2-1 bound more strongly to canonical NKX2-1 binding sites in AT1 cells (Little et al., 2021) following MEKi, but binding to HNF4α target sites in the duodenum (Chen et al., 2020) was attenuated (**Figure S6H**). In summary, we observed a striking degree of overlap between *Hnf4a* deletion and RAS/MEK inhibition in hybrid-ID LUAD and have shown that regulation of NKX2-1 chromatin binding is a likely mechanism by which KRAS inhibitors induce alveolar cell identity changes.

Since induction of AT1 identity has been proposed as a resistance mechanism to RAS inhibition (Li et al., 2024) and *Hnf4a* deletion also induces AT1 marker expression, we hypothesized that *Hnf4a* deletion may enhance resistance to MRTX1133. Bulk RNA-seq of organoids treated with a low dose of MRTX1133 +/- *Hnf4a* deletion revealed that the combination of *Hnf4a* deletion and KRAS inhibition led to a greater induction of AT1 marker genes than either modulation alone (**Figure S7A, Supplemental Table S9**). Surprisingly, however, *Hnf4a* deletion sensitized DP organoids to MRTX1133 rather than promoting resistance (**Figure 6J-K**). Cell death was also induced following *Hnf4a* deletion and KRAS inhibition to a greater extent than with either treatment alone (**Figure 6L-M**). Interestingly, deletion of *Foxa1/2* using *Foxa1/2* conditional DP organoids and cell lines also enhanced sensitivity to KRAS^G12D^ inhibition, likely reflecting the role of FoxA1/2 as upstream regulators of HNF4α and GI identity in hybrid-ID LUAD (**Figure S7B-C**). Together, our data show that RAS/MEK signaling and HNF4α coordinately induce a hybrid identity state within NKX2-1-positive LUAD, with important implications for response to targeted therapy (**Figure S7D**).

## DISCUSSION

LUAD tumor evolution is marked by a gradual loss of AT2 lineage fidelity and an epigenetic transition through alternate cell identity programs that enhance the malignant potential of tumor cells (LaFave et al., 2020; Marjanovic et al., 2020; Orstad et al., 2022; Tavernari et al., 2021; Yang et al., 2022). These profound identity changes culminate in hybrid-ID states whereby highly plastic tumor cells simultaneously activate divergent differentiation programs, preceding the eventual loss of both pulmonary and non-pulmonary fates to unleash poorly differentiated, metastasis-prone tumor cells. Here, we show that the GI lineage specifier, HNF4α, is a key driver of tumor growth and hybrid-ID differentiation in NKX2-1-positive LUAD. HNF4α coordinately induces GI/hepatic-like identity programs and dampens pulmonary programs, likely by physically interacting with NKX2-1 and altering its binding and activity. We also identified an unexpected role for RAS/MEK signaling in maintaining the hybrid-ID state in LUAD and uncovered a unifying mechanism – regulation of NKX2-1 chromatin binding – that underlies therapeutically relevant cell identity changes in response to both RAS/MEK targeted therapy and modulation of HNF4α.

HNF4α is critical for the growth of hybrid-ID LUAD. The growth promoting effects of GI/hepatic differentiation states within NKX2-1-positive LUAD likely stems from a combination of factors that will require additional investigation. First, several direct downstream targets of HNF4α are likely to be important for growth, including general metabolic enzymes such as IDH1 involved in key pro-tumor pathways (Calvert et al., 2017; Xu et al., 2020). Moreover, activation of tissue-specific metabolic enzymes by HNF4α may permit LUAD cells to utilize fuel sources that are typically unavailable to cells with a normal pulmonary identity. For example, metabolism of fructose as an energy source is a phenomenon typically restricted to a small subset of normal tissues including the liver and intestines (Herman & Birnbaum, 2021; Jang et al., 2018), but our data show that HNF4α activates enzymes associated with fructose metabolism (i.e. *SORD, ALDOB*) (**Supplemental Table S2**) with downstream effects on metabolite profiles (i.e. sorbitol levels, **Figure S3D**). Changes to cancer cell metabolism, including activation of enzymes involved in xenobiotic metabolism (**Figure S3C**), might also play a role in HNF4α-induced resistance to KRAS^G12D^ inhibition. Second, NKX2-1 restrains malignant progression of KRAS-mutant LUAD (Orstad et al., 2022; Snyder et al., 2013; Winslow et al., 2011). Thus, the ability of HNF4α to dampen NKX2-1 transcriptional activity likely also contributes to its tumorigenic effects. Finally, aberrant expression of HNF4α within a pulmonary identity appears to impair overall differentiation (e.g. **Figure 2E**), which likely promotes tumor progression. This observation is somewhat counterintuitive given that HNF4α evolved to impose specific identities in the GI tract and liver. We speculate that, in the absence of a normal GI/hepatic epigenome and/or proteome, HNF4α is not restricted to activation of a single identity. Moreover, co-expression of opposing lineage specifiers is sufficient to induce pluripotency in differentiated murine and human cells (Montserrat et al., 2013; Shu et al., 2013). This suggests that cancer cells might undergo dedifferentiation not only by progressive loss of individual identity programs, but also by the co-activation of opposing lineage specifiers. Importantly, evidence of hybrid identity states has been documented in other cancer types, including in pancreatic ductal adenocarcinoma (PDAC), where single cells simultaneously expressing markers of the two major PDAC identity states (classical and basal-like) have been characterized (Raghavan et al., 2021), suggesting that this might be a general mechanism of cancer progression.

Typically, lineage-specifying transcription factors guide cells along discrete developmental trajectories by activating tissue-specific gene expression programs. In contrast, here we find that NKX2-1 and HNF4α deviate from their specialized roles as lineage-defining transcription factors by physically interacting and binding to both GI/hepatic-like and pulmonary genes within hybrid-ID LUAD. The precise molecular implications of this novel protein complex in LUAD remain unclear. For instance, NKX2-1 and HNF4α may antagonize each other’s activity when these proteins are complexed, or they may simply sequester one another away from their respective canonical binding sites without direct functional impact. Additionally, gel filtration chromatography suggests that multiple NKX2-1 protein complexes exist in hybrid-ID LUAD, and the interactions between other cofactors, NKX2-1 binding partners, and HNF4α remain to be investigated. Our findings also suggest that transcriptional drivers of distinct identity programs might interact and/or interplay within hybrid-ID states in other cancer types.

Our data reveal that RAS/MEK signaling promotes the expression of GI/liver-like programs and HNF4α in NKX2-1-positive LUAD, highlighting this pathway as a key promoter of hybrid-ID differentiation. Moreover, the induction of HNF4α/GI-like programs via RAS/MEK signaling may be one mechanism by which increased RAS/MEK activity, downstream of oncogenic KRAS mutations, promotes LUAD tumor progression. Importantly, our results suggest that HNF4α and the gastric-like state contribute to intrinsic resistance to KRAS inhibition in NKX2-1-positive LUAD, suggesting that combinatorial therapeutic strategies targeting GI-like LUAD cells alongside KRAS inhibition could improve treatment outcomes. Recent studies in PDAC suggest that the classical subtype of PDAC, which is marked by high expression of HNF4α, is more resistant to KRAS inhibition than basal PDAC (Dilly et al., 2024; Singhal et al., 2024). This observation raises the intriguing possibility that HNF4α may also be a mediator or a biomarker of KRAS inhibitor resistance in cancer types beyond LUAD.

In addition to effects on GI/liver differentiation, we also found that RAS/MEK signaling is a critical regulator of NKX2-1 chromatin binding. This has general implications for LUAD treatment, as most targeted therapies inhibit this pathway and are therefore predicted to alter NKX2-1 activity. Moreover, this observation provides a likely molecular mechanism for the observation that KRAS inhibition induces therapeutically relevant alveolar identity programs in LUAD (Li et al., 2024), as these programs are largely NKX2-1 dependent (Little et al., 2019; Little et al., 2021; Snyder et al., 2013). Additionally, our observations that NKX2-1 binds more strongly to regions enriched for TEAD motifs following MEK inhibition raise the possibility that NKX2-1 might play a role in YAP/TEAD signaling-induced resistance to targeted therapy in LUAD (Edwards et al., 2023; Kurppa et al., 2020), in addition to effects on AT1/2 differentiation. Importantly, we observe HNF4α-independent changes to NKX2-1 localization, suggesting that this mechanism may have broad relevance across all NKX2-1-positive LUAD cases. The precise mechanisms by which RAS/MEK signaling alters NKX2-1 activity and/or localization remains to be determined. Notably, ERK can phosphorylate NKX2-1 and dampen its transcriptional activity by unclear mechanisms in vitro (Missero et al., 2000), suggesting that ERK may directly act on NKX2-1 within LUAD. In summary, our findings reveal novel mechanisms by which opposing lineage specifiers and oncogenic signaling pathways dictate pro-growth differentiation states and implicate HNF4α as a potential therapeutic target and/or biomarker of therapy resistance in LUAD.

## EXPERIMENTAL MODELS AND SUBJECT DETAILS

### Animal Studies

Mice harboring *Kras^FSF-G12D^* (Young et al., 2011), *Rosa-FSF-Cre^ERT2^* (Schonhuber et al., 2014), *Hnf4a*^flox^ (Hayhurst et al., 2001), and *p53^frt^* (Lee et al., 2012) have been previously described. All animals were maintained on a mixed 129/B6 background. All experimental mice were between 2 and 6 months of age at intubation. Mice of both sexes were used throughout each study, though the effect of sex on study results was not assessed. Animal studies were approved by the IACUC of the University of Utah, conducted in compliance with the Animal Welfare Act Regulations and other federal statutes relating to animals and experiments involving animals, and adhered to the principles set forth in the Guide for the Care and Use of Laboratory Animals, National Research Council (PHS assurance registration number A-3031-01).

### Husbandry and housing conditions of experimental animals

Experimental mice were mated or purchased. Appropriate housing conditions, including temperature, humidity, and light cycles, were maintained, and the mice were provided with a consistent diet. Transmission of transgenic and/or knockout alleles was monitored via DNA isolated from ear biopsy. Animals identified as negative for the presence of relevant alleles were used as control littermates or euthanized. Each mouse was marked by ear tagging and assigned a unique number. To preserve the genetic integrity of mouse models, careful handling practices were followed, and detailed records were kept throughout the duration of the study.

### Cell lines and primary cultures

All primary murine organoid cultures were established within Matrigel (Preclinical Research Shared Resource core facility) submerged in recombinant organoid medium for approximately two weeks (Advanced DMEM/F-12 supplemented with 1X B27 (Gibco), 1X N2 (Gibco), 1.25mM nAcetylcysteine (Sigma), 10mM Nicotinamide (Sigma), 10nM Gastrin (Sigma), 100ng/ml EGF (Peprotech), 100ng/ml R-spondin1 (Peprotech), 100ng/ml Noggin (Peprotech), and 100ng/ml FGF10 (Peprotech)). After organoids were established, cultures were switched to 50% L-WRN conditioned media (Miyoshi & Stappenbeck, 2013). Organoid lines were tested periodically for mycoplasma contamination. To maintain organoid cultures mycoplasma free, all culture media were supplemented with 2.5μg/ml Plasmocin.

3311 and HEK293T cells were cultured in DMEM/10% FBS (Gibco). NCI-H358, NCI-H441, and NCI-H2009 were cultured in RPMI/10% FBS (Gibco). NCI-H1651 was cultured in Advanced DMEM F-12/10% FBS/10mM HEPES (Gibco). All cell lines were tested periodically for mycoplasma contamination. To maintain cell cultures mycoplasma free, all culture media were supplemented with 2.5μg/ml Plasmocin.

## METHOD DETAILS

### Tumor initiation and tamoxifen administration in vivo

Autochthonous lung tumors were initiated by administering viruses via intratracheal intubation. Adenoviral mCMV-FlpO (4×10^7^ PFU/mouse) was used to initiate tumors for tumor burden and BrdU experiments. Adenoviral mSPC-FlpO (1×10^9^ PFU/mouse) was used to initiate tumors for scRNA-seq experiments. Adenoviruses were obtained from University of Iowa Viral Vector Core.

Tumor-specific activation of Cre^ERT2^ nuclear activity was achieved by intraperitoneal injection of tamoxifen (Sigma) dissolved in corn oil at a dose of 120mg/kg. Mice received 4 injections over the course of 5 days. BrdU incorporation was performed by injecting mice at 40mg/kg (Sigma) intraperitoneally 1 hour prior to tissue collection.

### Analysis of human lung adenocarcinoma

#### TCGA

We first filtered all TCGA Pan Cancer Atlas lung adenocarcinoma samples based on high NKX2-1 expression (*NKX2-1* gene expression z-score>-0.2, n=397/510 LUAD samples as of July 2024). The 397 NKX2-1-high samples were further filtered based on *HNF4A* expression. The *HNF4A*-high group was derived of patients with an *HNF4Α* gene expression z-score>0.8. Manual investigation of pathology reports was performed on all *HNF4A*-high patients, and patients with documented negative NKX2-1 (TTF-1) status were discarded. The final *HNF4A*-high (n=47) and *HNF4A*-low (n=339) groups are recorded in **Supplemental Table S1**. We identified differentially expressed genes between the *HNF4A*-high and *HNF4A*-low cohorts using DESeq2 (**Supplemental Table S2**) and used the resultant log2FCs to run GSEA-preranked analysis with MSigDB c8 gene sets (**Supplemental Table S3**). Signature scores for the curated HNF4α-regulated gene set were generated using ssGSEA. *EPS8L3*-high and low groups were identified by stratifying NKX2-1-positive patients by *EPS8L3* gene expression levels. The top 47 patients with highest expression of *EPS8L3* were selected as the *EPS8L3*-high group (**Supplemental Table S1**). Kaplan Meyer survival curves were generated using R. Survival analysis was performed using a Cox proportional hazards model. Differences in survival between groups is shown using hazard ratios and log-rank test p-values.

### Histology and immunohistochemistry

All tissues were fixed in 10% formalin overnight and when necessary, lungs were perfused with formalin via the trachea. Organoids were first fixed in 10% formalin overnight and then mounted in HistoGel (Thermo Fisher Scientific). Mounted organoids and tissues were transferred to 70% ethanol, embedded in paraffin, and four-micrometer sections were cut. Immunohistochemistry (IHC) was performed manually on Sequenza slide staining racks (Thermo Fisher Scientific). Sections were treated with Bloxall (Vector Labs) followed by Horse serum (Vector Labs) or Rodent Block M (Biocare Medical), primary antibody, and HRP-polymer-conjugated secondary antibody (anti-Rabbit from Vector Labs; anti-Mouse from Biocare). The slides were developed with Impact DAB (Vector Labs) and counterstained with hematoxylin. Slides were stained with antibodies against NKX2-1 (1:2000, Abcam, EP1584Y), HA-tag (1:400, CST, C29F4), or HNF4α (1:500, CST, C11F12). Images were taken on a Nikon Eclipse Ni-U microscope with a DS-Ri2 camera and NIS-Elements software. Histological analyses were performed on hematoxylin and eosin-stained and IHC-stained slides using NIS-Elements software. All histopathologic analysis was performed by a board-certified anatomic pathologist (E.L.S.).

### Establishing primary murine LUAD organoids

KPH mice bearing lung tumors were euthanized and lungs were isolated. Individual macroscopic tumors were removed from lungs, minced under sterile conditions, and digested at 37°C for 30 min with continuous agitation in a solution of Advanced DMEM/F12 containing the following enzymes: Collagenase Type I (Thermo Fisher Scientific, 450U/ml), Dispase (Corning, 5U/ml), DNaseI (Sigma, 0.25mg/ml). Enzymatic reactions were stopped by addition of cold Advanced DMEM/F-12 with 10% FBS. The digested tissue was repeatedly passed through a 20-gauge syringe needle, sequentially dispersed through 100μm, 70μm, and 40μm cell strainers, and treated with erythrocyte lysis buffer (eBioscience) to obtain a single cell suspension.

Organoid cultures were established by seeding 1×10^5^ tumor cells in 50μl of Matrigel (Corning) and plated in 24-well plates. Matrigel droplets were overlaid with recombinant organoid medium as described above (*Cell lines and primary cultures*). Two weeks after organoid establishment, cultures were switched to 50% L-WRN conditioned media. Organoid cultures were screened via immunohistochemistry and qPCR, and lines that expressed both HNF4α and NKX2-1 were selected for subsequent analysis. To generate uniform, HNF4α/NKX2-1-positive organoid cultures for downstream analysis, 3D organoid lines were subcloned as previously described (Pleguezuelos-Manzano et al., 2020) to generate 3154 and 1334 organoid lines. The full names of these organoid lines are SC3154#5 and SC1334#6 but names were shortened for clarity.

The 2D 3311 cell line and 1027B organoid line utilized in this study were derived as described in a previous study from our research group (Orstad et al., 2022). Notably, these cultures also contained conditional alleles of *Foxa1/2* but were not cultured in 4OHT and were phenotypically KP for these experiments.

### In vitro 4-hydroxytamoxifen (4OHT) treatment

Cells were transiently treated with 2mM 4OHT (Cayman Chemical Company, dissolved in 100% Ethanol) or vehicle (EtOH) for 48hr to activate Cre^ERT2^ nuclear activity and generate isogenic pairs of organoids.

### Generating a single cell suspension from organoid cultures

Matrigel domes containing organoids were broken down via repeated pipetting in Cell Recovery Solution (Corning, 500µl per Matrigel dome). Cell Recovery Solution containing organoids was transferred to sterile conical tubes and submerged in ice for 20-30min before centrifugation at 4°C (300-500G). Cell Recovery Solution supernatant was removed, and the cell pellet was washed in 1X PBS. Cells were then resuspended in pre-warmed TrypLE Express Enzyme (Thermo Fisher Scientific) and incubated for 10min at 37°C. TrypLE reaction was quenched via dilution with cold Splitting Media (Advanced DMEM/F-12 [Gibco], 10mM HEPES [Invitrogen], 1X Penicillin-Streptomycin-Glutamine [Invitrogen]). Cells were centrifuged and then resuspended in a pre-warmed DNase solution (L-WRN media supplemented to a final concentration of 200U/ml DNase [Worthington], 2.5mM MgCl2, 500mM CaCl2) and incubated for 10min at 37°C. Cells were centrifuged and washed in PBS before use.

### Immunoblotting

Protein was extracted by lysing cells on ice for 20min in RIPA buffer (50mM Tris HCl pH 7.4, 150mM NaCl, 0.1% (w/v) sodium dodecyl sulfate, 0.5% (w/v) sodium deoxycholate, 1% (v/v) Triton X-100) plus Pierce Protease Inhibitor (PPI, Thermo Fisher Scientific, A32959). Cellular debris was pelleted for 15min at 4°C and protein concentration was quantitated with the Pierce Bradford Protein Assay Kit (Thermo Fisher Scientific, 23200). A total of 15μg (organoids) or 30μg (cell lines) of protein lysates were separated on Tris-Glycine precast gels (SMOBIO, QP4510) and transferred to a nitrocellulose membrane (Thermo Fisher Scientific, 88018). Membranes were probed overnight with antibodies against FoxA1 (1:1000, Abcam 23738), FoxA2 (1:1000, Abcam 108422), NKX2-1 (1:2000, Abcam 133638), HNF4α (1:1000, CST C11F12), pERK (1:1000, CST 4370), total ERK (1:2000, CST 4695), and Vinculin (1:20000, Abcam 129002). The next day, membranes were probed with IRDye 800CW Goat anti-Rabbit IgG or IRDye 680RD Goat anti-Mouse IgG secondary antibodies (1:20000, LI-COR) and imaged with a LI-COR Odyssey CLx and Image Studio Software.

### Gel filtration chromatography

3311 KP cells (~7 x 10^7^) were collected and washed twice with PBS on ice. All of the following procedures were performed at 4°C. Washed cells were lysed in 2mL Farnham lysis buffer (5mM PIPES pH 8.0, 85mM KCl, 0.5% NP40) with Pierce Protease Inhibitor (PPI, Thermo Fisher Scientific, A32959) and incubated on ice for 10min. Cells were centrifuged at 1000 RCF for 5min at 4°C, and the pellet was resuspended in 2mL gel filtration lysis buffer with PPI (50mM Tris pH 8.0, 150mM NaCl, 1% Nonidet P-40 substitute, 10% glycerol) and incubated on ice for 30 min. The lysate was centrifuged at 3,500 rpm for 10min. The supernatant was further cleared by centrifugation at 35,000 rpm for 1 hour and then separated by size exclusion chromatography on a Superdex 200 column in gel filtration lysis buffer on an ÄKTA Pure 25L protein purification system. Samples were collected in fractions of 1.0mL. Immunoblotting was performed as described above.

### Lentiviral production and transduction

HEK293T cells were transfected with doxycycline-inducible TRE-HNF4α lentiviral vector (Camolotto et al., 2021), d8.9 packaging vector and VSV-G envelope vector mixed with TransIT-293 (Mirus Bio). Virus-containing supernatant was collected 48 and 72 hours after transfection, centrifuged to pellet floating HEK293T cells, and filtered using 0.45μm filters before storing long term at −80°C.

For stable transduction of organoids, organoid cultures were first prepared into single cell suspensions as described above (*Generating a single cell suspension from organoid cultures*). Cells were then resuspended in a 1:1 mixture, by volume, of 50% L-WRN and thawed lentivirus-containing supernatant. After supplementation with 8µg/ml polybrene, cells were incubated for 24 hours. Cells were then pelleted, mixed back with Matrigel, and seeded. 72 hours later, Blasticidin selection (10μg/mL) for 1 week was performed to achieve stable lines. Murine 3311 cells and human H1651 cells were transduced by culturing with undiluted lentiviral media containing 8μg/ml polybrene for 48 hours total, refreshing the media and polybrene at 24 hours. Cells were selected for 1 week with 12μg/mL Blasticidin. To induce HNF4α expression, cells and organoids were treated with 0.1-2.5μg/mL doxycycline for at least 48h.

### Subcutaneous cell line xenografts

For subcutaneous xenograft experiments, H1651 cells transduced with HNF4α lentiviral overexpression vector were collected and mixed with Matrigel 1:1 by volume. Cells were subcutaneously injected into the flanks of NOD/SCID-gamma chain deficient mice (NSG) (~3 million cells/flank). Mice were fed with normal chow or doxycycline-containing chow (Teklad Diet TD.01306) to induce HNF4α expression within implanted tumor cells. Tumor dimensions were measured with calipers, and tumor volume was calculated using the (L x W^2^)/2 formula. Tumor volume was monitored and measured every 7 days, and mice were euthanized once one tumor within the cohort surpassed 1000mm^3^.

### PrestoBlue Cell Viability Assays

Organoids were broken down into a single cell suspension as described above (*Generating a single cell suspension from organoid cultures*) and seeded at equal density with 5μl of Matrigel per well in a solid wall, clear bottom 96 well plate with 100μl of LWRN per well. For *Hnf4a* deletion growth experiments, cells were pretreated with 4OHT (or Ethanol) for 48 hours prior to seeding. For experiments involving drug treatments, organoids were given drug and/or 4OHT one day after seeding. A baseline measurement was taken one day after seeding by adding 10μl of PrestoBlue HS Cell Viability Reagent (Invitrogen) to each well and incubating at 37°C for 30min. After incubation, fluorescent emission was quantified on a Synergy HTX plate reader (Excitation: 528/20, Emission: 590/20). Presto blue reagent was removed, and wells were washed with warm PBS before adding fresh LWRN media. Measurements were taken every other day for *Hnf4a* deletion growth experiments or after 72 hours for drug treatment experiments.

### DAPI Cell Death Assay

For *Hnf4a* deletion experiments, 3154 and 1334 organoids were treated with 4OHT to delete *Hnf4a* for 7 days. For KRASi experiments, 3154 and 1334 organoids were treated with 4OHT to delete *Hnf4a* and/or 3nM MRTX1133 (Mirati Therapeutics) to inhibit KRAS^G12D^ for 72h. Organoids were harvested, broken down into a single cell suspension as described above (*Generating a single cell suspension from organoid cultures*), and resuspended in annexin V binding buffer (Invitrogen) to a density of ~2 x 10^5^ cells/mL. Cells were stained with DAPI (Sigma) and were immediately analyzed by flow cytometry on a BD Fortessa flow cytometer to identify DAPI-positive (dead) and DAPI-negative (live) cells.

### EdU Cell Proliferation Assay

3154 and 1334 DP organoids were treated with 4OHT (or ethanol) for 48h. The following protocol was adapted from the Click-iT EdU Alexa Fluor Flow Cytometry Assay Kit (Invitrogen). Organoids were treated for 30 minutes with 0.01mM EdU in normal LWRN media. Organoids were immediately collected and broken down into a single cell suspension as described above (*Generating a single cell suspension from organoid cultures*). Organoids were washed in PBS with 1% BSA and were fixed in 4% formaldehyde for 15 minutes, before washing twice more with PBS + 1% BSA. Just before analysis by flow cytometry, cells were permeabilized in 1X Saponin (Invitrogen) in PBS/BSA, and 500μL of a Click-iT cocktail was added to each sample (100mM CuSO_4_, 500mM Ascorbic Acid, and 5mM Alexa Fluor 642 fluorescent azide dye (Invitrogen)). Cells were incubated for 30 minutes protected from light, washed in 1X Saponin in PBS/BSA and immediately analyzed by flow cytometry on a BD Fortessa flow cytometer for EdU positive cells.

### Proximity Ligation Assay

2D cell lines (3311 or H1651) were grown on glass coverslips to ~80% confluency. For HNF4α overexpression 3311 cells, cells were treated with 2.5μg/mL doxycycline for 72hr. Cells were washed twice in cold PBS, fixed in cold methanol at −20°C for 15mins, and washed twice more in cold PBS. Duolink PLA (Sigma) protocol was then followed according to the manufacturer’s specifications using Anti-Rabbit PLUS (Sigma, DUO92002) and Anti-Mouse MINUS (Sigma, DUO92004) probes and green detection reagent kit (Sigma, DUO92014). The following antibodies and concentrations were utilized: NKX2-1 (1:2000, Abcam, ab76013), HNF4α (1:1000, Perseus Proteomics, PP-H1415), FoxA1 Rabbit (1:4000, Abcam, ab173287), FoxA1 Mouse (1:1000, Millipore, 2F83). Coverslips were mounted onto microscopy slides using mountant containing DAPI (Abcam, ab104139). Slides were guarded from light and imaged within 48 hours on a confocal microscope (Leica SP8 Confocal White Light). Average PLA signals per nucleus was quantified using ImageJ software (Fiji Version 2.14.0) for a minimum of 3 fields of view per condition.

### RNA extraction, cDNA synthesis, and qPCR

RNA was isolated via Trizol-chloroform extraction followed by column-based purification. The aqueous phase was brought to a final concentration of 35% ethanol, and RNA was purified using the PureLink RNA Mini Kit (Thermo Fisher Scientific) according to the manufacturer’s specifications.

cDNA was synthesized from Trizol-extracted RNA using LunaScript RT SuperMix (NEB, M3010) according to manufacturer’s specifications. qPCR was performed on cDNA using Luna Universal Probe qPCR Master Mix (NEB, M3004) and Taqman probes (Thermo Fisher) according to manufacturer’s specifications, and 35 cycles were used for the denaturation and extension steps. Transcript levels were normalized to PPIA and quantitated by the ΔΔCt method.

### RNA sequencing

RNA was collected from 2 biological replicates of the following conditions: 3154 KP/KPH isogenic organoid cultures (2 weeks following 4OHT treatment), H1651 cells overexpressing HNF4α (7 days of treatment with 1μg/mL doxycycline), 3311 MEKi treated cells (48h of treatment with 50nM Trametinib (LC Laboratories, T-8123)), 1027B MEKi or KRASi treated organoids (48h of treatment with 10nM Trametinib or 20nM MRTX1133 (Mirati Therapeutics)), H1651 MEKi treated cells (48h of treatment with 20nM Trametinib), H358 MEKi treated cells (48h of treatment with 20nM Trametinib), and 3154 organoids +/- HNF4α +/- MRTX1133 (48h of treatment with 3nM MRTX1133 (Mirati Therapeutics) +/- 4OHT). Cells were collected directly into Trizol and stored at −80°C until purification. For organoids, 4 confluent 20μL Matrigel domes were collected per sample.

RNA was isolated via Trizol-chloroform extraction followed by column-based purification. The aqueous phase was brought to a final concentration of 35% ethanol, and RNA was purified using the PureLink RNA Mini kit according to the manufacturer’s instructions (Thermo Fisher Scientific). Library preparation was performed using the NEBNext Ultra II Directional RNA Library Prep with poly(A) mRNA isolation. Sequencing was performed using the Illumina NovaSeq 6000 (150 x 150 bp paired-end sequencing, 25 million reads per sample).

### RNA-seq data processing and analysis

The mouse mm10 (or mm39 for 3154 MRTX1133+/-*Hnf4a* experiment) or human hg38 genome and gene feature files were downloaded from Ensembl and a reference database was created using STAR version 2.7.6a (Dobin et al., 2013). Optical duplicates were removed from fastq files using Clumpify v38.34 (Bushnell, 2021). Reads were trimmed of adapters and aligned to the reference database using STAR in two-pass mode to output a BAM file sorted by coordinates. Mapped reads were assigned to annotated genes using featureCounts v1.6.3 in paired-end mode (Liao et al., 2019). Raw counts were filtered to remove features with zero counts and features with five or fewer reads in every sample. DEGs were identified using the hciR package (https://github.com/HuntsmanCancerInstitute/hciR) with a 5% false discovery rate and DESeq2 version 1.42.1 (Love et al., 2014). GSEA-Preranked was run with the differential gene list generated from DESeq2 and the following gene sets: c5, c8 and Hallmarks gene sets from MSigDB and custom gene sets found in **Supplemental Table S3**. Gene sets smaller than 15 and larger than 500 were excluded from analysis.

### GC-MS metabolomics

3154 organoids were treated with ethanol (vehicle) or 4OHT to delete *Hnf4a* for 6 days. Organoids were broken down into a single cell suspension and biological replicates of 5 x 10^6^ cells were flash frozen and subjected to metabolite extraction and analysis.

### Metabolite Extraction

The following protocol was performed at the Metabolomics Core Facility at the University of Utah. To each sample was added cold 90% methanol (MeOH) solution to give a final concentration of 80% MeOH to each cell pellet. Samples were then incubated at −20°C for 1 hr. After incubation the samples were centrifuged at 20,000 x g for 10 minutes at 4°C. The supernatant was then transferred from each sample tube into a labeled, fresh micro centrifuge tube. Pooled quality control samples were made by removing a fraction of collected supernatant from each sample and process blanks were made using only extraction solvent and no cell culture. The samples were then dried *en vacuo*.

### Mass Spectrometry Analysis of Samples

All GC-MS analysis was performed with an Agilent 5977b GC-MS MSD-HES and an Agilent 7693A automatic liquid sampler. Dried samples were suspended in 40µL of a 40mg/mL O-methoxylamine hydrochloride (MOX) (MP Bio #155405) in dry pyridine (EMD Millipore #PX2012-7) and incubated for one hour at 37°C in a sand bath. 25µL of this solution was added to auto sampler vials. 60µL of N-methyl-N-trimethylsilyltrifluoracetamide (MSTFA with 1% TMCS, Thermo #TS48913) was added automatically via the auto sampler and incubated for 30 minutes at 37°C. After incubation, samples were vortexed and 1µL of the prepared sample was injected into the gas chromatograph inlet in the split mode with the inlet temperature held at 250°C. A 10:1 split ratio was used for analysis for the majority of metabolites. Any metabolites that saturated the instrument at the 10:1 split were analyzed at a 50:1 split ratio. The gas chromatograph had an initial temperature of 60°C for one minute followed by a 10°C/min ramp to 325°C and a hold time of 10 minutes. A 30-meter Agilent Zorbax DB-5MS with 10 m Duraguard capillary column was employed for chromatographic separation. Helium was used as the carrier gas at a rate of 1 mL/min.

### Analysis of Mass Spectrometry Data

Data was collected using MassHunter software (Agilent). Metabolites were identified and their peak area was recorded using MassHunter Quant. Metabolite identity was established using a combination of an in-house metabolite library developed using pure purchased standards, the NIST library and the Fiehn library.

### LC-MS Lipidomics

3154 organoids were treated with ethanol (vehicle) or 4OHT to delete *Hnf4a* for 6 days. Organoids were broken down into a single cell suspension and biological replicates of 1 x 10^6^ cells were flash frozen and subjected to lipid extraction and analysis.

### Lipidomics Sample Preparation

The following protocol was performed at the Metabolomics Core Facility at the University of Utah. Lipids were extracted using the Matyash method, a biphasic solvent system of Methyl tert-Butyl Ether (MTBE), Methanol (MeOH), and water (Matyash et al., 2008) with some modifications. Cell culture samples had 213µL of PBS added first, vortexed briefly, and 25µL aliquots taken out for future protein assay. In a randomized order lipids were then extracted from Eppendorf’s by adding 975µL of MeOH and MTBE solution containing internal standards (Avanti EquiSPLASH LIPIDOMIX at 5µL per sample, Cholesterol-d7 (79µg/mL) 5µL per sample, Palmitic Acid-d31 (100µg/mL) 10µL per sample). Samples rested on ice for 60 minutes with brief vortexes every 15 minutes, and then were centrifuged at 4°C for 10 minutes at 14,000xg with the upper phase being collected. Another 1mL aliquot of solution consisting of MTBE/MeOH/H_2_O (10: 3: 2.5) was added for re-extraction, followed by a brief vortex and set to rest at room temperature for 15 minutes. The samples were then centrifuged again at 4°C for 10 minutes at 14,000xg where after the upper phases would be combined and set to evaporate till dryness under speedvac. Once samples dried they were re-suspended in 1000µL of (4:1:1) IPA/ACN/H_2_O, vortexed, and centrifuged at 4°C for 10 minutes at 14,000xg. The extracts were then placed into LCMS vials with inserts for analysis; concurrently, a process blank sample was produced along with a pooled quality control (QC).

### Mass Spectrometry Analysis of Samples

Lipid extracts were separated on an Acquity UPLC CSH C18 column (2.1 x 100 mm; 1.7 µm) coupled to an Acquity UPLC CSH C18 VanGuard precolumn (5 × 2.1 mm; 1.7 µm) (Waters, Milford, MA) maintained at 65°C connected to an Agilent HiP 1290 Sampler, Agilent 1290 Infinity pump, and Agilent 6545 Accurate Mass Q-TOF dual AJS-ESI mass spectrometer (Agilent Technologies, Santa Clara, CA). Samples were analyzed in a randomized order in both positive and negative ionization modes in separate experiments acquiring with the scan range m/z 100 – 1700. For positive mode, the source gas temperature was set to 225 °C, with a drying gas flow of 11 L/min, nebulizer pressure of 40 psig, sheath gas temp of 350 °C and sheath gas flow of 11 L/min. VCap voltage is set at 3500 V, nozzle voltage 500V, fragmentor at 110 V, skimmer at 85 V and octopole RF peak at 750 V. For negative mode, the source gas temperature was set to 300 °C, with a drying gas flow of 11 L/min, a nebulizer pressure of 30 psig, sheath gas temp of 350 °C and sheath gas flow 11 L/min. VCap voltage was set at 3500 V, nozzle voltage 75 V, fragmentor at 175 V, skimmer at 75 V and octopole RF peak at 750 V. Mobile phase A consisted of ACN:H_2_O (60:40, *v/v*) in 10mM ammonium formate and 0.1% formic acid, and mobile phase B consisted of IPA:ACN:H_2_O (90:9:1, *v/v/v*) in 10mM ammonium formate and 0.1% formic acid. For negative mode analysis the modifiers were changed to 10mM ammonium acetate. The chromatography gradient for both positive and negative modes started at 15% mobile phase B then increased to 30% B over 2.4 min, it then increased to 48% B from 2.4 – 3.0 min, then increased to 82% B from 3 – 13.2 min, then increased to 99% B from 13.2 – 13.8 min where it’s held until 16.7 min and then returned to the initial conditions and equilibrated for 5 min. Flow was 0.4 mL/min throughout, with injection volumes of 2µL for positive and 8µL negative mode. Tandem mass spectrometry was conducted using iterative exclusion, the same LC gradient at collision energies of 20 V and 27.5 V in positive and negative modes, respectively.

### Analysis of Mass Spectrometry Data

Data processing was performed using Agilent MassHunter (MH) Workstation and software packages MH Qualitative and MH Quantitative. The pooled QC (n=8) and process blank (n=4) were injected throughout the sample queue to ensure the reliability of acquired lipidomics data. For lipid annotation, accurate mass and MS/MS spectral matching was used with LipidMatch library (Koelmel et al., 2017). Results from the positive and negative ionization modes from Lipid Annotator were merged based on the class of lipid identified. Data exported from MH Quantitative was evaluated using Excel where initial lipid targets are parsed based on the following criteria. Only lipids with relative standard deviations (RSD) less than 30% in QC samples are used for data analysis. Additionally, only lipids with background AUC counts in process blanks that are less than 30% of QC are used for data analysis. The parsed excel data tables are normalized based on the ratio to class-specific internal standards prior to statistical analysis.

### Statistical Analysis and Data Visualization for Metabolomics and Lipidomics

Multivariate analysis was performed using MetaboAnalyst (Xia et al., 2015).Statistical models were created for the normalized data after logarithmic transformation (base 10) and Pareto scaling. Heatmaps were generated in R using normalized abundance values from MetaboAnalyst analysis. Raw and processed metabolomics and lipidomics data can be found in **Supplemental Table S7**.

### Single-cell RNA sequencing

KPH mice were injected with tamoxifen or corn oil 10 weeks after intubation, and two weeks later, single cells were collected as follows. Lungs and heart were perfused with PBS. Individual macroscopic tumors were removed from the lungs and broken down into single cell suspensions as described in *Establishing primary murine LUAD organoids* methods. Pre-depletion cells were viably cryopreserved in 5% DMSO/FBS. The remaining cells were depleted of CD45-positive and CD31-positive cells using MACS with Miltenyi microbeads (CD45:130-052-301; CD31:130-097-418) and LD columns (130-042-901) following manufacturer recommendations. Post-depletion cells were viably cryopreserved in 5% DMSO/FBS. The day of library preparation, cells were thawed, washed in cold PBS, and resuspended in PBS + 0.5% BSA. Cells were stained with DAPI and sorted for single, live cells using a BD FACSAria cell sorter.

Protocols used to generate scRNA-seq data with 10x Genomics Chromium platform can be found at https://support.10xgenomics.com/single-cell-gene-expression. In brief, the Chromium Single Cell Gene Expression Solution with 3’ chemistry, version 3 (PN-1000075) was used to barcode individual cells with 16bp 10X barcodes and to tag cell specific transcript molecules with 10bp Unique Molecular Identifier (UMI) according to the manufacturer’s instructions. The following protocol was performed at the High-Throughput Genomics Shared Resource at Huntsman Cancer Institute, University of Utah. Single cells were suspended in PBS with 0.04% BSA, and the cell suspension was passed through a 40 micron cell strainer. Viability and cell count were assessed on Countess II (Thermo Fisher Scientific). Suspensions were equilibrated to targeted cell recovery of 10,000 cells. 10x Gel Beads and reverse transcription reagents were added and cell suspensions were loaded to Chromium Single Cell A (PN-120236) to form Gel Beads-in emulsions (GEMs) - the nano-droplets. Within individual GEMs, cDNA generated from captured and barcoded mRNA was synthesized by reverse transcription at the setting of 53°C for 45min followed by 85°C for 5min. Subsequent A tailing, end repair, adaptor ligation and sample indexing were performed in bulk according to the manufacturer’s instructions. The resulting barcoding libraries were qualified on Agilent D1000 ScreenTape on Agilent Technology 2200 TapeStation system and quantified by quantification PCR using KAPA Biosystems Library Quantification Kit for Illumina Platforms (KK4842). Multiple libraries were then normalized and sequenced on NovaSeq 6000 with 2 × 150 PE mode.

### scRNAseq data processing and analysis

#### Demultiplexing and data alignment

Single-cell RNA-seq data from both KP (n=2) and KPH (n=2) tumors were demultiplexed using the 10x cellranger mkfastq version 7.2.0 to create fastq files with the I1 sample index, R1 cell barcode+UMI, and R2 sequence. Reads were aligned to the mouse genome (mm10 with custom CRE-ERT2 reference) and UMIs were generated using cellranger count 7.2.0 with include-introns set to true. For the KP sample without stromal depletion, we captured 7,658 cells total with 13,600 mean molecules detected per cell and 2,967 median genes per cell. For the KP sample with stromal depletion, we captured 7,284 cells total with 19,906 mean molecules detected per cell and 4,416 median genes per cell. For the KPH sample without stromal depletion, we captured 6,558 cells total with 14,127 mean molecules detected per cell and 3,032 median genes per cell. For the KPH sample with stromal depletion, we captured 7,867 cells total with 18,017 mean molecules detected per cell and 4,283 median genes per cell. Additional details of the primary Cell Ranger data processing can be found at: https://support.10xgenomics.com/single-cell-gene-expression/software/pipelines/latest/algorithms/overview.

#### Quality control, clustering, and cell type identification

Single cell expression data was subjected to common Seurat V4 workflows for initial quality control and clustering (Hao et al., 2021) (https://satijalab.org/seurat/articles/pbmc3k_tutorial.html). Cells with unique feature counts over 7500 or less than 500 and over 12% mitochondrial counts were filtered out for downstream analysis. Counts of cells passing QC were then log normalized and scaled based on all genes using Seurat’s NormalizeData and ScaleData functions. PCA linear dimension reduction was performed and Seurat’s FindNeighbors function was employed to embed single cell profiles in a K-nearest neighbor (KNN) graph based on a (30 PC) PCA space. The FindClusters function was utilized to iteratively group cells together using Louvian algorithm modularity optimization techniques. Clustering was performed based on the top 30 dimensions using Seurat’s RunUMAP function. Differentially expressed genes for each cluster were identified using Seurat’s FindMarkers function using default parameters. Following QC filtering and tumor cell identification, 6823 KP and 7544 KPH high quality tumor cells remained. For subsequent analyses, identified tumor cells from KP and KPH tumors were subsetted out and reclustered based on the top 12 dimensions. Cell barcodes identified as tumor cells that were used for downstream analyses are included in **Supplemental Table S4.** Reclustering of identified tumor cells in UMAP space revealed 11 clusters (**Supplemental Table S4**) To identify KPH complete recombinants, expression of the floxed fourth and fifth exons of *Hnf4a* were quantified (**Supplemental Table S4**).

#### Differential gene expression and signature score assignment

Differentially expressed genes in each of the tumor cell clusters were calculated using Seurat’s FindMarkers function. Differentially expressed genes of UMAP clusters from all KP and KPH tumor cells can be found in **Supplemental Table S4**. Gene module scores for published gene signatures for gastric-like LUAD, AT2-like LUAD and AT1 cells (Du et al., 2015; Marjanovic et al., 2020), were determined per cell across the tumor cell dataset using Seurat’s AddModuleScore function. Gene lists for each of these scores can be found in **Supplemental Table S3**. CytoTRACE scores were calculated using the CytoTRACE2 R Package v1.0.0 ( https://github.com/digitalcytometry/cytotrace2) (Gulati et al., 2020) with default parameters to report a continuous score ranging from 0 (most differentiated) to 1 (least differentiated).

#### Data imputation

We used a data imputation approach to model the expression of low-detection genes in our scRNA-seq dataset (Linderman et al., 2022). The ALRA method utilizes a low-rank approximation approach to impute gene expression values while preserving biological zeros.

### ChIP-sequencing

For *Hnf4a* deletion ChIP-seq, 3154 organoids were treated with 4OHT (or ethanol) for 6 days to delete *Hnf4a*. For H1651 ChIP-seq, H1651 cells harboring a lentiviral HNF4α overexpression construct were treated with 1μg/mL doxycycline for 7 days. For MEKi ChIP-seq, 3311 cells were treated with 50nM Trametinib (LC Laboratories, T-8123) for 48h, and 1027B organoids and H358 cells were treated with 20nM Trametinib for 48h to inhibit the MAPK pathway. All ChIP-seq experiments were performed in biological duplicates. For cell line ChIP-seq, ~1 x 10^7^ cells were collected per ChIP experiment. For organoid ChIP-seq, ~4 x 10^6^ cells were collected per ChIP experiment. For organoid ChIP experiments, organoids were first collected in Cell Recovery Solution and incubated on ice for 30mins. Organoids were then washed three times in cold PBS. On the second wash, PBS was supplemented with DNase solution (containing a final concentration of 200U/ml DNase [Worthington], 2.5mM MgCl2, 500mM CaCl2). 2D adherent cell lines were washed 2X in cold PBS, then cells were scraped and collected and washed one more time. Cell/organoid pellets were then resuspended in 5ml of 2mM DSG buffer (1X PBS, 1 mM MgCl_2_) containing 40μL of 0.25M disuccinimidyl glutarate (DSG) (Thermo Fisher Scientific) and rotated at room temperature for 35 min. Methanol-free formaldehyde was then added to a final concentration of 1% and cells were crosslinked for 10min. The cross-linking reaction was quenched with the addition of glycine to a final concentration of 125mM. Cells were washed with cold PBS, then the cell pellet was snap frozen in liquid nitrogen and stored at −80°C.

Cell pellets were thawed on ice for 5 minutes then lysed in 1mL of Farnham lysis buffer (5mM PIPES pH 8.0, 85mM KCl, 0.5% NP40) with Pierce Protease Inhibitor (PPI, Thermo Fisher Scientific, A32959). Samples were centrifuged at 4°C (1000G) then resuspended in 1mL of RIPA lysis buffer (1X PBS, 1% NP40, 0.5% sodium deoxycholate, 0.1% sodium dodecyl sulfate) with PPI. Chromatin was sonicated with a QSonica Q800R (pulse: 30 s on/30 s off; sonication time: 20 minutes; amplitude: 70%). After sonication, samples were centrifuged at 17,000 x g for 15 minutes and an input was collected from each sample of sheared chromatin. Chromatin from each sample was then immunoprecipitated overnight with 5μg of the following antibodies (per sample) premixed with Protein G Dynabeads (for mouse antibodies; Thermo Fisher Scientific, 10004D) or Protein A Dynabeads (for rabbit antibodies; Thermo Fisher Scientific, 10002D): HNF4α (Perseus Proteomics PPH1415), NKX2-1 (Abcam ab133737), FoxA1 (Abcam, ab170933), FoxA2 (CST 8186). The next day, bound chromatin was washed 5 times with LiCl Wash Buffer (100mM Tris pH 7.5, 500mM LiCl, 1% NP-40, 1% sodium deoxycholate) and crosslinks were reversed by incubating samples in IP Elution Buffer (1% SDS, 0.1% NaHCO_3_) overnight at 65°C. ChIP DNA was purified using Zymo ChIP DNA Clean and Concentrator Kit (Zymo, D5205).

Library preparation was performed using the ChIP-seq with NEBNext DNA Ultra II library prep kit using Unique Molecular Indexes (UMIs). Sequencing was performed using the Illumina NovaSeq 6000 (150 x 150 bp paired-end sequencing, 25 million reads per sample).

### ChIP-seq data processing and analysis

Fastq alignments were pre-processed with the merge_umi_fastq application from the UMIScripts package (https://github.com/HuntsmanCancerInstitute/UMIScripts) to associate the UMI sequence, provided as a third Fastq file, into the read comment. Reads were aligned using Bowtie2 v2.2.9 (Langmead & Salzberg, 2012) to the standard chromosomes of the mouse genome (version mm10) or the human genome (version hg38). Duplicate alignments based on the UMI code were removed using the bam_umi_dedup application (UMIScripts) allowing for 1 mismatch. Peaks were called using MACS2 v2.2.7 (Zhang et al., 2008) with a significance of q-value < 0.01. Coverage tracks were generated with MACS2 as Reads Per Million. Input libraries were obtained from all cell line and organoid samples and were used as controls for each ChIP-seq experiment. All ChIP-seq experiments were performed in biological duplicates. Peaks called in both biological replicates were identified using Bedtools v2.28.0 (Quinlan & Hall, 2010) with a 1-bp minimum overlap to generate a consensus list of peaks for downstream analysis. Genomic annotation of binding sites was performed using HOMER (Heinz et al., 2010). Motif analysis was performed on 100-bp regions surrounding the summit of identified peaks using the HOMER suite. Peaks were annotated to genes using the annotatePeaks.pl function from the HOMER suite to annotate each peak to the closest TSS. Publicly available ChIP-seq data was intersected with the ChIP-seq datasets generated in this study using Bedtools with bed files downloaded from published data (Chen et al., 2019; Gu et al., 2024; Little et al., 2021; Qu et al., 2021). Integrative Genomics Browser (IGV) (Robinson et al., 2011) was utilized to visualize ChIP-seq peaks.

Differential ChIP-seq peaks were identified using the Diffbind package v3.4.11 (https://bioconductor.org/packages/release/bioc/vignettes/DiffBind/inst/doc/DiffBind.pdf) with a q-value cutoff < 0.05 using edgeR (Robinson et al., 2010) for the analysis method. Motif finding for differential ChIP-seq peaks was performed using HOMER using the entire 400bp differential peak region output by DiffBind. Heat maps and profile plots were generated using deeptools v3.5.1 (Ramirez et al., 2016). ATAC-seq coverage files from different cell states in the KP mouse model were downloaded from (LaFave et al., 2020) and ATAC-seq signal at differentially bound NKX2-1 peaks following *Hnf4a* deletion was visualized using deeptools. Pathway analysis was performed on annotated differential ChIP-seq peaks using Enrichr (Chen et al., 2013).

### ChIP-re-ChIP

We optimized a sequential ChIP (ChIP-re-ChIP) protocol in part using a published experimental workflow (Seneviratne et al., 2024). For ChIP-re-ChIP experiments, 3311 cells (~30 million per ChIP) were collected, sonicated, and immunoprecipitated as described above (*ChIP-sequencing*) with the addition of a pre-clearing step where lysates were incubated with protein A and G dynabeads prior to immunoprecipitation for 1hr at 4°C. Samples were immunoprecipitated with antibodies against FoxA1 or HNF4α after removing an aliquot of chromatin for input controls. After washing with LiCl Wash Buffer, beads were resuspended in 10mM DTT and incubated with agitation at 37°C for 40 minutes. DTT was diluted by adding 1mL RIPA buffer with PPI, and diluted chromatin was immunoprecipitated for a second time overnight with the reciprocal antibody (either FoxA1 or HNF4α). The remainder of the ChIP protocol and analysis was carried out as described above. HNF4α:FoxA1 and FoxA1:HNF4α ChIP-re-ChIP data was compared to single ChIP experiments carried out on the same cell line for NKX2-1, FoxA1, and HNF4α.

## QUANTIFICATION AND STATISTICAL ANALYSIS

All graphing and statistical analysis was performed with PRISM software or R, with all graphs showing mean and standard deviation or standard error as indicated in figure legends. The statistical details can be found in the corresponding figure legend. All NGS statistical analysis was performed according to published pipeline protocols cited, with a statistical significance cutoff of padj<0.05.

## ACKNOWLEDGEMENTS

We are grateful to members of the Snyder lab for helpful suggestions and comments. We thank T. Jacks and C. Cabana for helpful suggestions, B. Dalley for sequencing expertise, T. Parnell and J. Gertz for ChIP-seq expertise, J. Marvin for FACS expertise, and B. Spike and O. Allen for scRNA-seq expertise. The results published here are in part based upon data generated by the TCGA Research Network: https://www.cancer.gov/tcga. E.L.S. was supported by grants from the NIH (R01CA212415, R01CA240317 and R01CA237404), the American Lung Association (LCD-821670), and institutional funds (Department of Pathology and Huntsman Cancer Institute/Huntsman Cancer Foundation, University of Utah). G.F. was supported by the NIH/NCI (F31CA275328). G.F. was also supported by a Genetics Training Grant (T32GM141848). The graphical abstract for this publication was created with Biorender.com. Research reported in this publication utilized shared resources (including High Throughput Genomics, Bioinformatics, Flow Cytometry, Biorepository and Molecular Pathology, and The Center for High Performance Computing) at the University of Utah and was supported by the National Cancer Institute of the National Institutes of Health under award number P30CA042014. Work in the flow cytometry core was also supported by the National Center for Research Resources of the National Institutes of Health under Award Number 1S20RR026802-1. Metabolomics analysis was performed at the Metabolomics Core Facility at the University of Utah. Mass spectrometry equipment was obtained through NCRR Shared Instrumentation Grant 1S10OD016232-01, 1S10OD018210-01A1 and 1S10OD021505-01. The content is solely the responsibility of the authors and does not necessarily represent the official views of the NIH.

## AUTHOR CONTRIBUTIONS

G.F. and E.L.S. designed experiments. G.F., H.A., S.C., R.T., A.W., and K.T.O. performed experiments. G.F., H.A., and E.L.S. analyzed data. E.L.S. performed histopathologic review. G.F. and E.L.S. wrote the manuscript. All authors discussed results, reviewed, and revised the manuscript.

## RESOURCE AVAILABILITY

### Lead contact

Further information and requests for resources and reagents should be directed to and will be fulfilled by the lead contact, Eric Snyder

### Materials availability

Murine organoid lines and cell lines generated in this study are available upon request.

### Data and code availability

- Bulk RNA-seq (GSE281963), scRNA-seq (GSE281964), and ChIP-seq (GSE281961) data have been deposited at Gene Expression Omnibus and are available as of the date of publication.
- Microscopy data reported in this paper will be shared by the lead contact upon request.
- This paper does not report original code.
- Any additional information required to reanalyze the data reported in this work paper is available from the Lead Contact upon request.

## SUPPLEMENTAL INFORMATION

**Supplemental Figures S1-S7.**

**Table S1.** TCGA PanCancer Atlas *NKX2-1*-high LUAD samples, *HNF4A*-high and *HNF4A*-low groups and baseline characteristics, related to Figures 1, S1, 2, S2, S3, and 3.

**Table S2.** Raw counts, normalized counts and DEGs in KP vs KPH 3154 murine organoids, control vs HNF4α OE H1651 human cells, and *HNF4A*-low vs *HNF4A*-high TCGA Pan Cancer Atlas LUAD samples, related to Figures 2, S2, S3, 3, and S5.

**Table S3.** Curated gene set list and enriched gene sets by GSEA in KP vs KPH 3154 murine organoids, control vs HNF4α OE H1651 human cells, and *HNF4A*-low vs *HNF4A*-high TCGA Pan Cancer Atlas LUAD samples, related to Figures 2, S2, S3, and 3.

**Table S4.** scRNA-seq metadata, cluster associations, DEGs per cluster, HNF4α exon counts, and DEGs and pathway analysis between selected clusters, related to Figures 2, S2, and 3.

**Table S5.** ChIP-seq peaks and gene annotations for KP and KPH 3154 murine organoid ChIP-seq, human H1651 cell line ChIP-seq, murine 3311 cell line control and MEKi ChIP-seq, murine 1027B organoid control and MEKi ChIP-seq, and human H358 cell line control and MEKi ChIP-seq, related to Figures 2, S2, 4, S4, 5, S5, 6, and S6.

**Table S6.** Differentially bound NKX2-1 peaks and gene annotations for KP vs KPH 3154 murine NKX2-1 ChIP-seq, 1027B organoid MEKi NKX2-1 ChIP-seq, and H358 cell line MEKi NKX2-1 ChIP-seq, related to Figures 5, S5, 6, and S6.

**Table S7.** Raw Metaboanalyst input, processed normalized/scaled abundance, and PCA scores from 3154 KP and KPH organoid GC-MS metabolomics and LC-MS lipidomics, related to Figure S3.

**Table S8.** ChIP-seq peaks and gene annotations for HNF4α:FoxA1 ChIP-re-ChIP experiments, related to Figures 4 and S4.

**Table S9.** Raw counts, normalized counts and DEGs in control vs RAS/MEKi 3311 murine cells, 1027B murine organoids, H358 cells, H1651 cells, and 3154 murine organoids +/-*Hnf4a* +/- KRASi, related to Figures 6, S6, and S7.

**Table S10.** Enriched gene sets by GSEA in control vs MEKi 3311 murine cells and 1027B murine organoids, related to Figure 6 and S6.

