## Supplemental Figures for "Opposing lineage specifiers induce a pro-tumor hybrid-identity state in lung adenocarcinoma"

### FIGURE S1

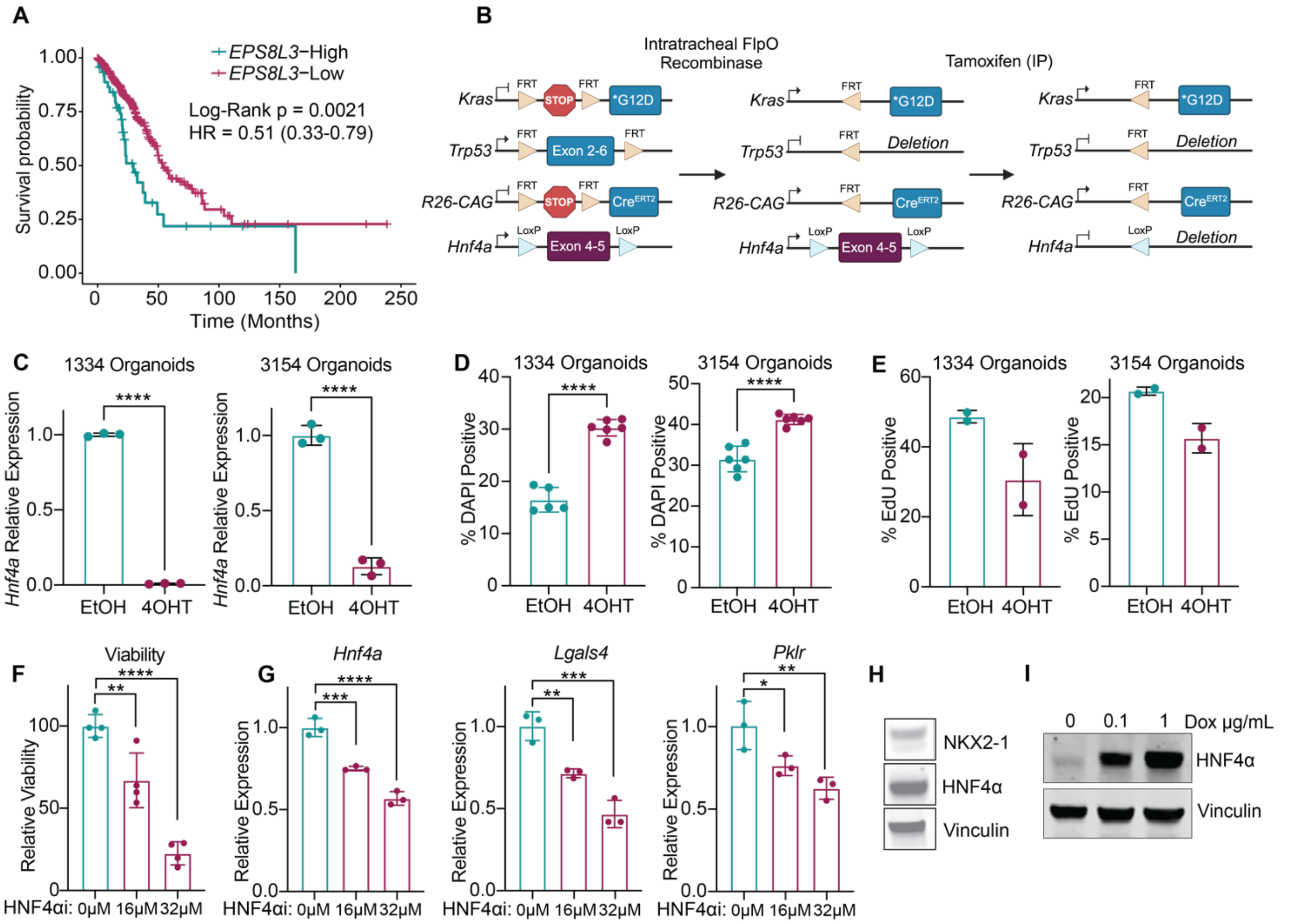
A
EPS8L3−High
EPS8L3−Low
Log-Rank p = 0.0021
HR = 0.51 (0.33-0.79)
Survival probability
Time (Months)
B
Intratracheal FlpO Recombinase
Tamoxifen (IP)
Kras
Trp53
R26-CAG
Hnf4a
FRT
LoxP
STOP
*G12D
Exon 2-6
Exon 4-5
Deletion
C
1334 Organoids
3154 Organoids
Hnf4a Relative Expression
EtOH
4OHT
D
% DAPI Positive
E
% EdU Positive
F
Viability
Relative Viability
HNF4αi: 0μM 16μM 32μM
G
Hnf4a
Lgals4
Pklr
Relative Expression
H
NKX2-1
HNF4α
Vinculin
I
0 0.1 1 Dox μg/mL
HNF4α
Vinculin

**Supplemental Figure 1. Related to Figure 1**.

**A.** Kaplan-Meier curve of PanCancer LUAD TCGA samples stratified based on high expression of *NKX2-1* and high (n=47) or low (n=339) expression of *EPS8L3*, excluding mucinous tumors. Log-Rank test and CoxPH hazard ratio with 95% CI shown.

**B.** Diagram of KPH mouse model utilized in this study.

**C.** qRT-PCR for *Hnf4a* mRNA expression in KPH organoids treated with EtOH or 4OHT to delete *Hnf4a* (one representative replicate shown of n=3 biological replicates; unpaired t test p<0.0001; error bars, SD).

**D.** DAPI staining for cell death in KPH organoids, measured by flow cytometry 7 days following treatment with EtOH (KP) or 4OHT (KPH) (n=2 biological replicates; unpaired t test ****p<0.0001; error bars, SD).

**E.** EdU incorporation measured by flow cytometry 2 days following treatment of KPH organoids with EtOH or 4OHT (n=2 biological replicates, error bars, SD).

**F.** Presto blue viability assay of DP organoids (3154) treated with 16 or 32μM BI6015 to inhibit HNF4α (one representative replicate shown of n=2 biological replicates; one-way ANOVA with Dunnett's multiple comparisons test, **p<0.005, ****p<0.0001; error bars, SD).

**G.** qRT-PCR for HNF4α downstream targets *Lgals4* and *Pklr* following inhibition of HNF4α with 16 or 32μM BI6015 for 72 hours in 3154 organoids (one representative replicate shown of n=2 biological replicates; one-way ANOVA with Dunnett's multiple comparisons test, *p<0.05,**p<0.006, ***p<0.0006,****p<0.0001; error bars, SD).

**H**. Representative immunoblot demonstrating co-expression of NKX2-1 and HNF4α in H1651 human LUAD cell line.

**I.** Representative immunoblot from H1651 cell line transduced with lentiviral dox-inducible overexpression construct utilized for subcutaneous tumor experiment demonstrating HNF4α overexpression in vitro pre-implantation. Cells were given 0.1 or 1μg/mL doxycycline for 3 days.

#### FIGURE S2

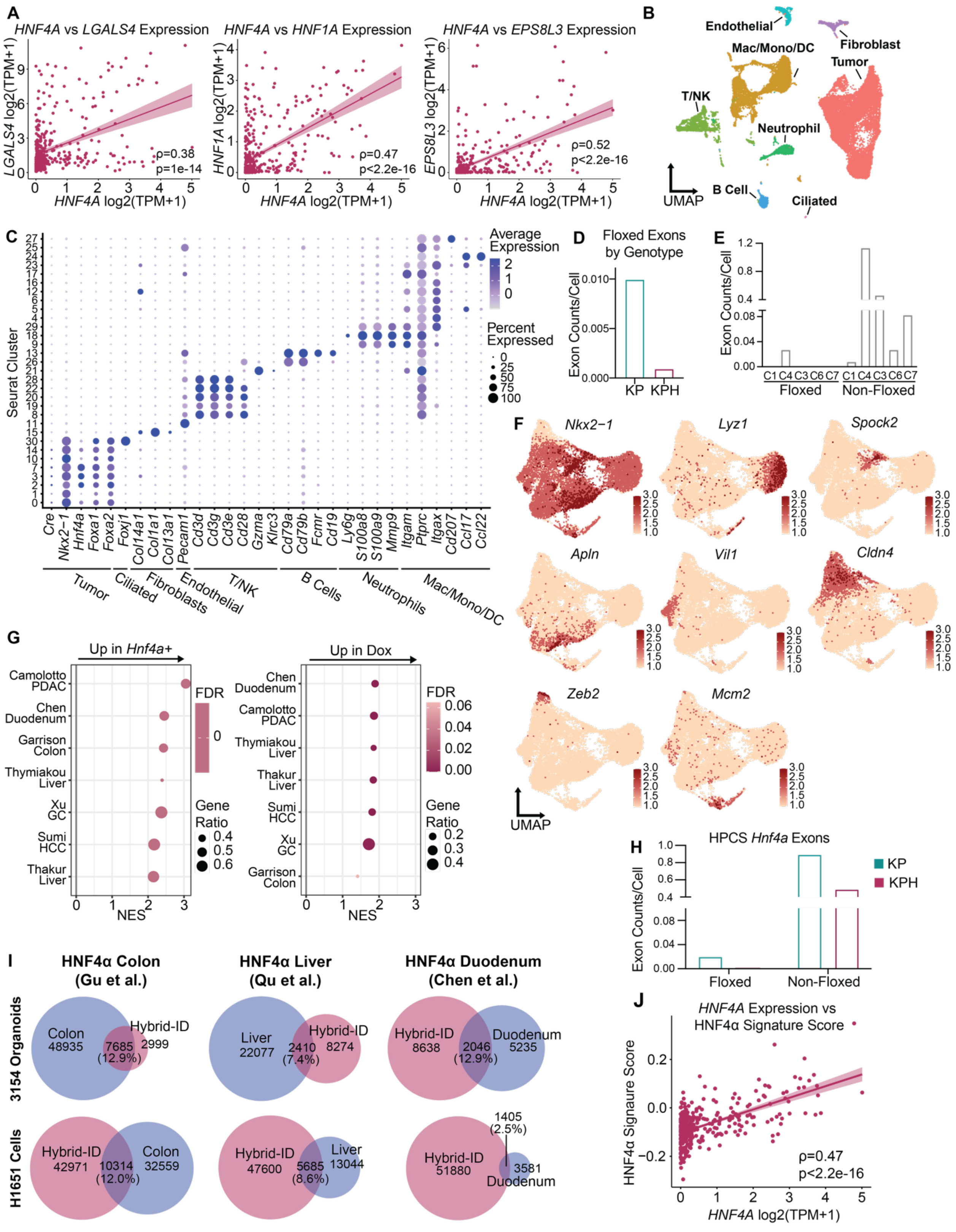
A
HNF4A vs LGALS4 Expression
HNF4A vs HNF1A Expression
HNF4A vs EPS8L3 Expression
ρ=0.38
p=1e-14
ρ=0.47
p<2.2e-16
ρ=0.52
p<2.2e-16
HNF4A log2(TPM+1)
LGALS4 log2(TPM+1)
HNF1A log2(TPM+1)
EPS8L3 log2(TPM+1)
B
Endothelial
Fibroblast
Mac/Mono/DC
Tumor
T/NK
Neutrophil
B Cell
Ciliated
UMAP
C
Average Expression
Percent Expressed
Seurat Cluster
Tumor
Ciliated
Fibroblasts
Endothelial
T/NK
B Cells
Neutrophils
Mac/Mono/DC
D
Floxed Exons by Genotype
Exon Counts/Cell
KP
KPH
E
Exon Counts/Cell
Floxed
Non-Floxed
F
Nkx2−1
Lyz1
Spock2
Apln
Vil1
Cldn4
Zeb2
Mcm2
UMAP
G
Up in Hnf4a+
Up in Dox
FDR
Gene Ratio
NES
Camolotto PDAC
Chen Duodenum
Garrison Colon
Thymiakou Liver
Xu GC
Sumi HCC
Thakur Liver
H
HPCS Hnf4a Exons
Exon Counts/Cell
KP
KPH
Floxed
Non-Floxed
I
HNF4α Colon (Gu et al.)
HNF4α Liver (Qu et al.)
HNF4α Duodenum (Chen et al.)
3154 Organoids
H1651 Cells
Colon 48935
Hybrid-ID 2999
7685 (12.9%)
Liver 22077
Hybrid-ID 8274
2410 (7.4%)
Hybrid-ID 8638
Duodenum 5235
2046 (12.9%)
Hybrid-ID 42971
Colon 32559
10314 (12.0%)
Hybrid-ID 47600
Liver 13044
5685 (8.6%)
Hybrid-ID 51880
1405 (2.5%)
3581 Duodenum
J
HNF4A Expression vs HNF4α Signature Score
HNF4α Signature Score
HNF4A log2(TPM+1)
ρ=0.47
p<2.2e-16

**Supplemental Figure 2. Related to Figure 2**.

**A.** Scatterplots of *HNF4A* expression vs expression of *LGALS4*, *HNF1A*, or *EPS8L3* (log2(TPM+1)) in curated TCGA LUAD cohort. Spearman correlation coefficients and p-values shown.

**B.** UMAP of high-quality KP (n=13,345) and KPH (n=12,835) cells colored by Seurat cluster and annotated by cell type.

**C.** Dot plot showing relative expression of cell type markers per Seurat cluster.

**D.** Number of reads per cell aligning to *Hnf4a* floxed exons 4 and 5 in KP vs KPH tumor cells.

**E.** Number of reads per cell aligning to *Hnf4a* floxed exons 4 and 5 vs non-floxed exons of *Hnf4a* in selected Seurat clusters.

**F.** UMAP of additional markers of identity programs within NKX2-1-positive LUAD: *Nkx2-1*, *Lyz1* (AT2), *Spock2* (AT1), *Apln* (Early Gastric), *Vil1* (Gastric), *Cldn4* (High Plasticity Cell State (HPCS)), *Zeb2* (EMT), *Mcm2* (G2M).

**G.** GSEA pathway analysis for published HNF4α-regulated gene signatures on differentially expressed genes in 3154 KPH murine organoid line (left) and H1651 dox-inducible HNF4α-overexpression cell line (right).

**H.** Number of reads per cell aligning to *Hnf4a* floxed exons in KP vs KPH cells within the HPCS cluster 2.

**I.** Quantification of overlapping peaks (number of overlapping peaks and percent overlap out of total peaks) between HNF4α ChIP-seq datasets from 3154 organoids and H1651 cells and publicly available HNF4α ChIP-seq datasets from normal murine liver, duodenum, and colon.

**J.** Scatterplot of ssGSEA enrichment scores for curated HNF4α regulated gene set across *NKX2-1*-positive TCGA samples vs expression of *HNF4A* (log2(TPM+1)). Spearman correlation coefficient and p-values shown.

### FIGURE S3

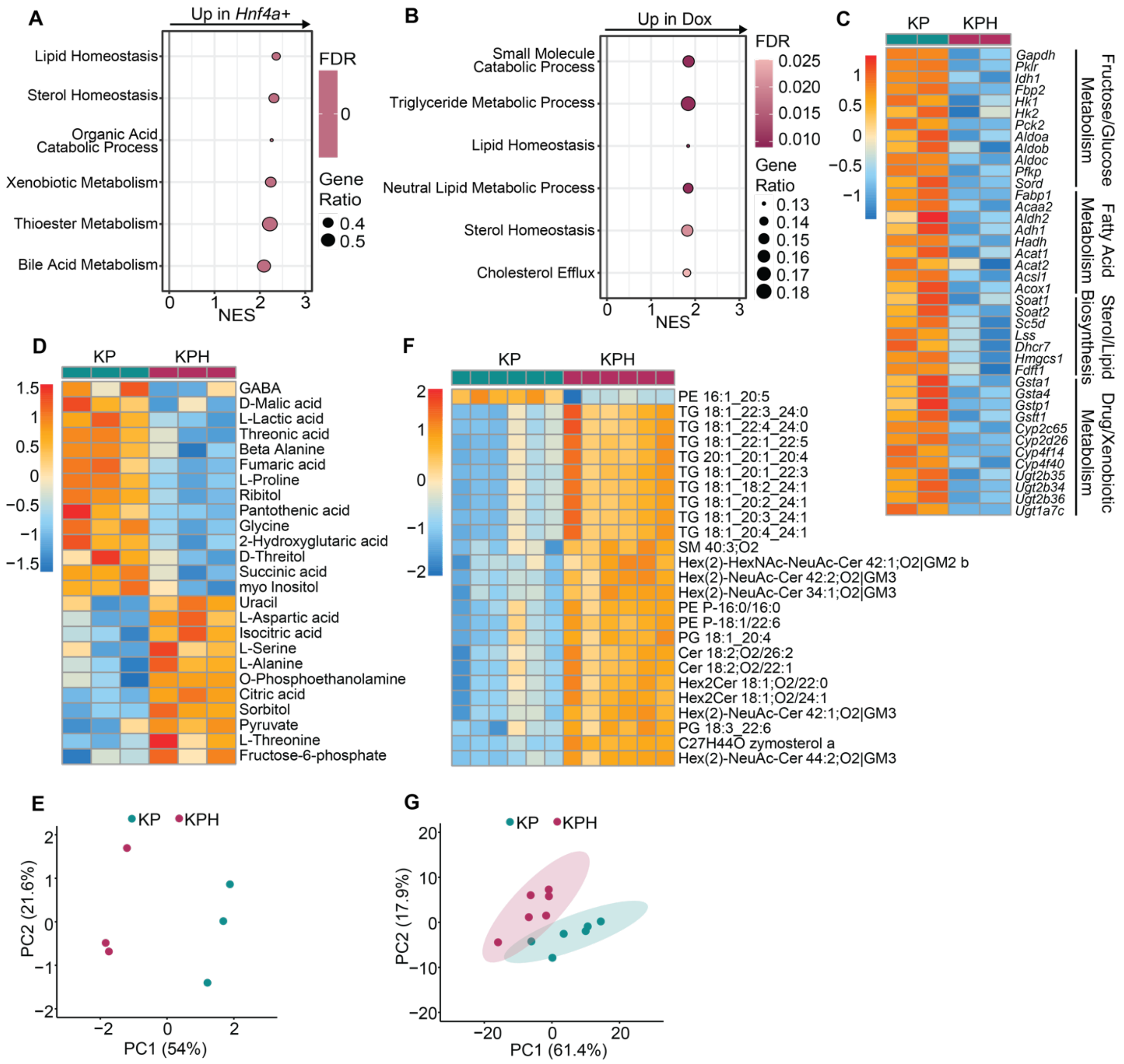
A
Up in Hnf4a+
Lipid Homeostasis
Sterol Homeostasis
Organic Acid Catabolic Process
Xenobiotic Metabolism
Thioester Metabolism
Bile Acid Metabolism
NES
FDR
Gene Ratio
B
Up in Dox
Small Molecule Catabolic Process
Triglyceride Metabolic Process
Lipid Homeostasis
Neutral Lipid Metabolic Process
Sterol Homeostasis
Cholesterol Efflux
NES
FDR
Gene Ratio
C
KP
KPH
Gapdh
Pklr
Idh1
Fbp2
Hk1
Hk2
Pck2
Aldoa
Aldob
Aldoc
Pfkp
Sord
Fabp1
Acaa2
Aldh2
Adh1
Hadh
Acat1
Acat2
Acsl1
Acox1
Soat1
Soat2
Sc5d
Lss
Dhcr7
Hmgcs1
Fdft1
Gsta1
Gsta4
Gstp1
Gstt1
Cyp2c65
Cyp2d26
Cyp4f14
Cyp4f40
Ugt2b35
Ugt2b34
Ugt2b36
Ugt1a7c
Fructose/Glucose Metabolism
Fatty Acid Metabolism
Sterol/Lipid Biosynthesis
Drug/Xenobiotic Metabolism
D
KP
KPH
GABA
D-Malic acid
L-Lactic acid
Threonic acid
Beta Alanine
Fumaric acid
L-Proline
Ribitol
Pantothenic acid
Glycine
2-Hydroxyglutaric acid
D-Threitol
Succinic acid
myo Inositol
Uracil
L-Aspartic acid
Isocitric acid
L-Serine
L-Alanine
O-Phosphoethanolamine
Citric acid
Sorbitol
Pyruvate
L-Threonine
Fructose-6-phosphate
F
KP
KPH
PE 16:1_20:5
TG 18:1_22:3_24:0
TG 18:1_22:4_24:0
TG 18:1_22:1_22:5
TG 20:1_20:1_20:4
TG 18:1_20:1_22:3
TG 18:1_18:2_24:1
TG 18:1_20:2_24:1
TG 18:1_20:3_24:1
TG 18:1_20:4_24:1
SM 40:3;O2
Hex(2)-HexNAc-NeuAc-Cer 42:1;O2|GM2 b
Hex(2)-NeuAc-Cer 42:2;O2|GM3
Hex(2)-NeuAc-Cer 34:1;O2|GM3
PE P-16:0/16:0
PE P-18:1/22:6
PG 18:1_20:4
Cer 18:2;O2/26:2
Cer 18:2;O2/22:1
Hex2Cer 18:1;O2/22:0
Hex2Cer 18:1;O2/24:1
Hex(2)-NeuAc-Cer 42:1;O2|GM3
PG 18:3_22:6
C27H44O zymosterol a
Hex(2)-NeuAc-Cer 44:2;O2|GM3
E
KP
KPH
PC2 (21.6%)
PC1 (54%)
G
KP
KPH
PC2 (17.9%)
PC1 (61.4%)

**Supplemental Figure 3**. **Related to Figure 2.**

**A.** GSEA pathway analysis on differentially expressed genes in control vs *Hnf4a*-deleted 3154 DP organoid line for representative metabolism-related gene signatures.

**B.** GSEA pathway analysis on differentially expressed genes in dox-inducible HNF4α-overexpression H1651 cell line for representative metabolism-related gene signatures.

**C.** Heatmap showing expression of selected metabolism-associated genes in control vs *Hnf4a*-deleted 3154 organoids. Z-scores across samples of log2 normalized counts shown.

**D.** Heatmap showing normalized abundance of top 25 differential metabolites upon deletion of *Hnf4a* in 3154 DP organoids. Row normalized by Z-score (one representative replicate shown of n=2 biological replicates).

**E.** PCA plot of metabolomic profiles of control and *Hnf4a*-deleted 3154 organoids (one representative replicate shown of n=2 biological replicates).

**F.** Heatmap showing normalized abundance of differential lipid species following deletion of *Hnf4a* in 3154 DP organoids. Row normalized by Z-score.

**G.** PCA plot of lipidomic profiles of 3154 control and *Hnf4a*-deleted 3154 organoids.

FIGURE S4

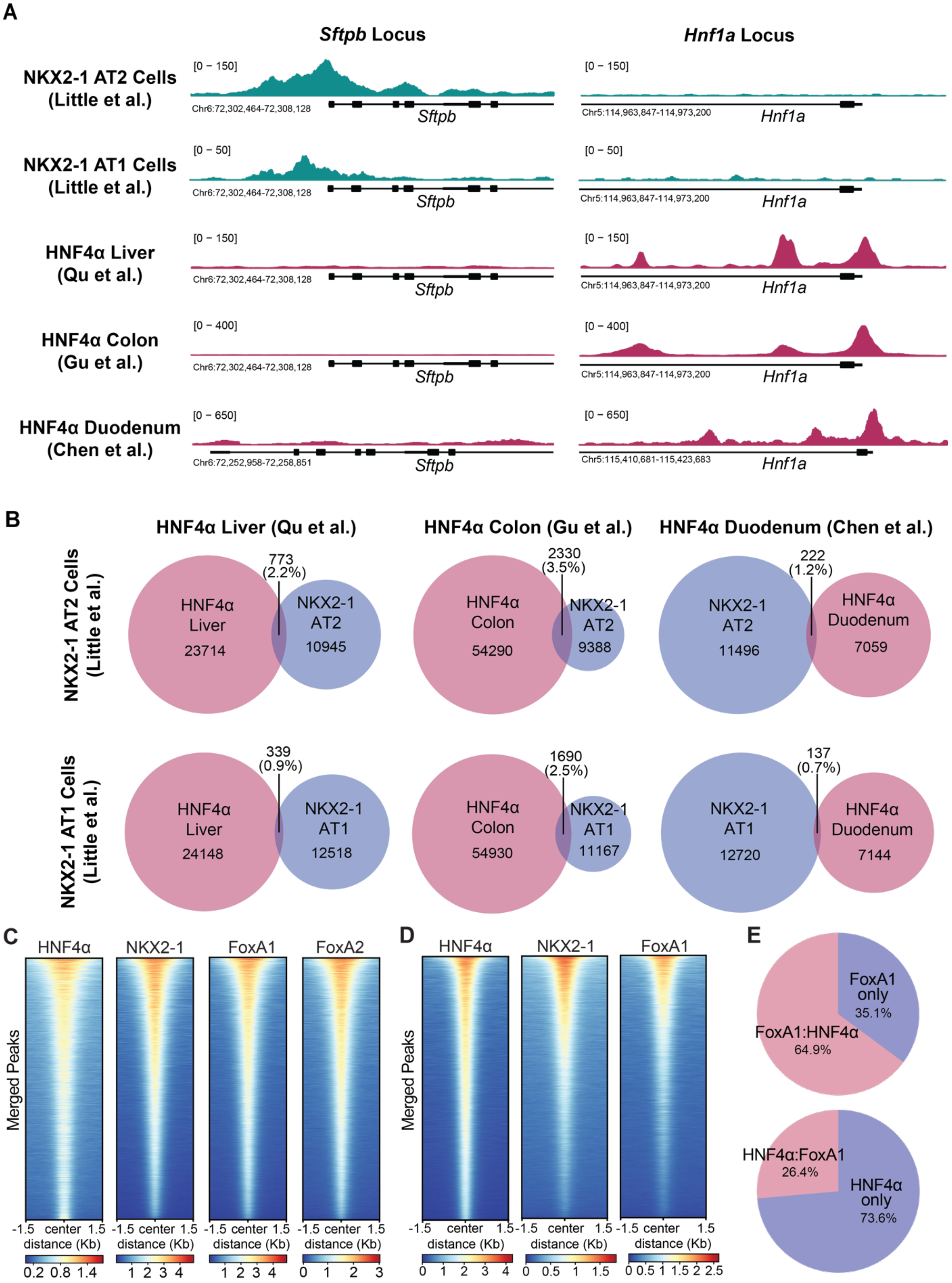
A
Sftpb Locus
Hnf1a Locus
NKX2-1 AT2 Cells (Little et al.)
NKX2-1 AT1 Cells (Little et al.)
HNF4α Liver (Qu et al.)
HNF4α Colon (Gu et al.)
HNF4α Duodenum (Chen et al.)
[0 – 150]
[0 – 50]
[0 – 400]
[0 – 650]
Chr6:72,302,464-72,308,128
Chr6:72,252,958-72,258,851
Chr5:114,963,847-114,973,200
Chr5:115,410,681-115,423,683
Sftpb
Hnf1a
B
HNF4α Liver (Qu et al.)
HNF4α Colon (Gu et al.)
HNF4α Duodenum (Chen et al.)
NKX2-1 AT2 Cells (Little et al.)
HNF4α Liver 23714
773 (2.2%)
NKX2-1 AT2 10945
HNF4α Colon 54290
2330 (3.5%)
NKX2-1 AT2 9388
NKX2-1 AT2 11496
222 (1.2%)
HNF4α Duodenum 7059
NKX2-1 AT1 Cells (Little et al.)
HNF4α Liver 24148
339 (0.9%)
NKX2-1 AT1 12518
HNF4α Colon 54930
1690 (2.5%)
NKX2-1 AT1 11167
NKX2-1 AT1 12720
137 (0.7%)
HNF4α Duodenum 7144
C
HNF4α
NKX2-1
FoxA1
FoxA2
Merged Peaks
-1.5 center 1.5
distance (Kb)
0.2 0.8 1.4
1 2 3 4
1 2 3 4
0 1 2 3
D
HNF4α
NKX2-1
FoxA1
Merged Peaks
-1.5 center 1.5
distance (Kb)
0 1 2 3 4
0 0.5 1 1.5
0 0.5 1 1.5 2 2.5
E
FoxA1 only 35.1%
FoxA1:HNF4α 64.9%
HNF4α:FoxA1 26.4%
HNF4α only 73.6%

**Supplemental Figure 4. Related to Figure 4**.

**A.** ChIP-seq tracks of published NKX2-1 and HNF4α ChIP-seq datasets from normal murine tissues at pulmonary and GI genes.

**B.** Quantification of overlapping peaks (number of overlapping peaks and percent overlap out of total peaks) between HNF4α ChIP-seq datasets from normal murine liver, duodenum, and colon and NKX2-1 ChIP-seq datasets from normal murine AT1 and AT2 cells.

**C.** Heatmap showing occupancy of designated transcription factors at merged peak regions (significant peaks called by Macs2 in at least 1 condition) from NKX2-1, HNF4α, FoxA1 and FoxA2 ChIP-seq experiments in 3154 DP murine organoid. Peaks ordered by descending mean signal across all datasets.

**D.** Heatmap showing occupancy of designated transcription factors at merged peak regions (significant peaks called by Macs2 in at least 1 condition) from NKX2-1, HNF4α and FoxA1 ChIP-seq experiments in H1651 human cell line. Peaks ordered by descending mean signal across all datasets.

**E.** Overlap between FoxA1 (top) and HNF4α (bottom) ChIP-seq peaks in 3311 cells and designated ChIP-re-ChIP peaks.

**FIGURE S5**

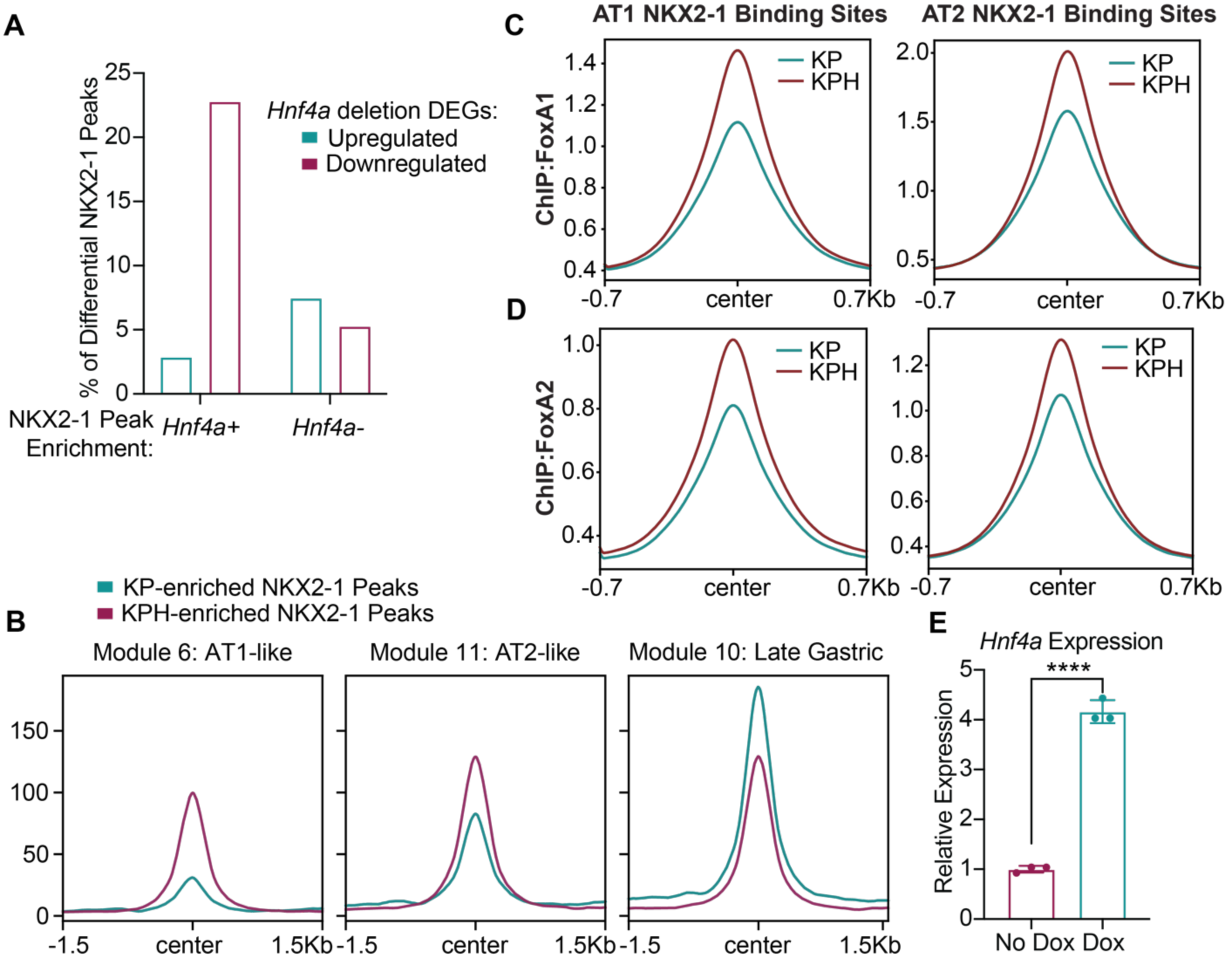
A
% of Differential NKX2-1 Peaks
Hnf4a deletion DEGs:
Upregulated
Downregulated
NKX2-1 Peak Enrichment:
Hnf4a+
Hnf4a-
B
KP-enriched NKX2-1 Peaks
KPH-enriched NKX2-1 Peaks
Module 6: AT1-like
Module 11: AT2-like
Module 10: Late Gastric
-1.5
center
1.5Kb
C
AT1 NKX2-1 Binding Sites
AT2 NKX2-1 Binding Sites
ChIP:FoxA1
KP
KPH
-0.7
center
0.7Kb
D
ChIP:FoxA2
E
Hnf4a Expression
Relative Expression
****
No Dox
Dox

**Supplemental Figure 5. Related to Figure 5**.

**A.** Association between annotated differential NKX2-1 binding sites in 3154 control vs *Hnf4a*-deleted organoids and differentially expressed genes from 3154 *Hnf4a* deletion bulk RNA-seq dataset.

**B.** Mean ATAC-seq signal from LaFave et al. scATAC-seq dataset at differential NKX2-1 peaks in AT1-like, AT2-like and Late Gastric-like clusters.

**C.** Mean peak profile of 3154 control and *Hnf4a*-deleted FoxA1 ChIP-seq signal at AT1-specific and AT2-specific NKX2-1 binding sites.

**D.** Mean peak profile of 3154 control and *Hnf4a*-deleted FoxA2 ChIP-seq signal at AT1-specific and AT2-specific NKX2-1 binding sites.

**E.** qRT-PCR showing overexpression of *Hnf4a* in 3311 cell line utilized for PLA experiment (one representative replicate shown of n=2 biological replicates; unpaired t test $p<0.0001$; error bars, SD).

### FIGURE S6

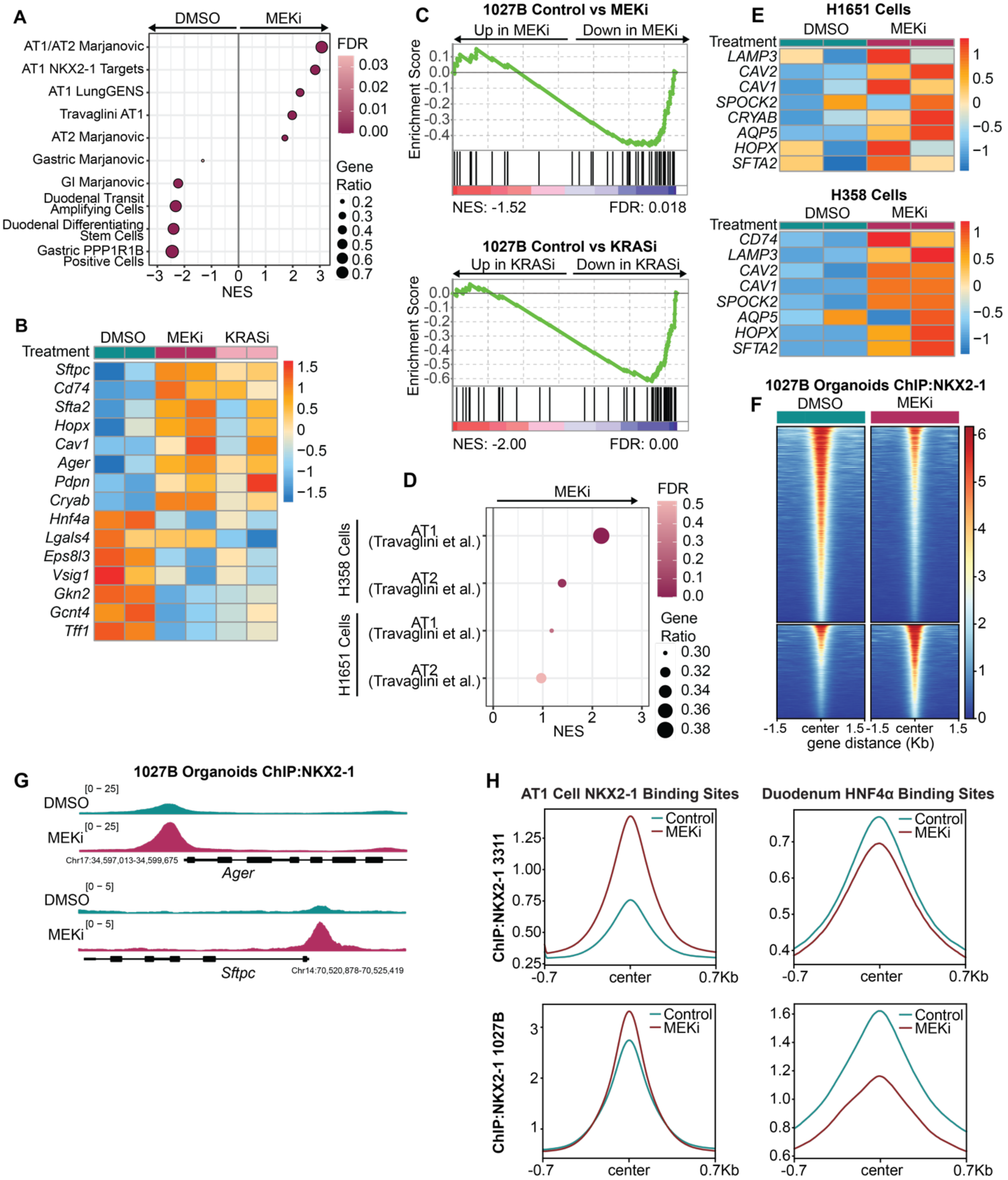
A
DMSO
MEKi
AT1/AT2 Marjanovic
AT1 NKX2-1 Targets
AT1 LungGENS
Travaglini AT1
AT2 Marjanovic
Gastric Marjanovic
GI Marjanovic
Duodenal Transit Amplifying Cells
Duodenal Differentiating Stem Cells
Gastric PPP1R1B Positive Cells
NES
FDR
Gene Ratio
B
Treatment
DMSO
MEKi
KRASi
Sftpc
Cd74
Sfta2
Hopx
Cav1
Ager
Pdpn
Cryab
Hnf4a
Lgals4
Eps8l3
Vsig1
Gkn2
Gcnt4
Tff1
C
1027B Control vs MEKi
Up in MEKi
Down in MEKi
Enrichment Score
NES: -1.52
FDR: 0.018
1027B Control vs KRASi
Up in KRASi
Down in KRASi
NES: -2.00
FDR: 0.00
D
MEKi
H358 Cells
H1651 Cells
AT1 (Travaglini et al.)
AT2 (Travaglini et al.)
NES
FDR
Gene Ratio
E
H1651 Cells
Treatment
DMSO
MEKi
LAMP3
CAV2
CAV1
SPOCK2
CRYAB
AQP5
HOPX
SFTA2
H358 Cells
CD74
F
1027B Organoids ChIP:NKX2-1
DMSO
MEKi
gene distance (Kb)
G
1027B Organoids ChIP:NKX2-1
DMSO
MEKi
[0 - 25]
[0 - 5]
Chr17:34,597,013-34,599,675
Ager
Sftpc
Chr14:70,520,878-70,525,419
H
AT1 Cell NKX2-1 Binding Sites
Duodenum HNF4α Binding Sites
Control
MEKi
ChIP:NKX2-1 3311
ChIP:NKX2-1 1027B
-0.7
center
0.7Kb

**Supplemental Figure 6. Related to Figure 6**.

**A.** GSEA pathway analysis on differentially expressed genes in control vs MEKi-treated (10nM Trametinib) DP organoid line (1027B). Representative cell type gene signatures shown.

**B.** Heatmap showing expression of selected gastrointestinal and pulmonary marker genes in control and MEKi or $KRAS^{G12D}$ inhibitor-treated DP 1027B organoids. Z-scores across samples of log2 normalized counts shown.

**C.** GSEA enrichment plots for curated HNF4α-regulated gene set in 1027B DP organoids treated with Trametinib (MEKi) or MRTX1133 (KRASi).

**D.** GSEA enrichment anlaysis for AT1 and AT2 gene signatures in human NKX2-1-positive LUAD cell lines treated with Trametinib (MEKi).

**E.** Heatmap showing expression of selected pulmonary marker genes in control and MEKi-treated human cell lines. Z-scores across samples of log2 normalized counts shown.

**F.** Heatmap depicting NKX2-1 occupancy at differential NKX2-1 binding sites following MEKi in 1027B organoids. Differential binding analysis was performed using DiffBind with an adjusted p value cutoff of <0.05 which identified 7577 control-specific peaks (out of 40,415 total control NKX2-1 peaks) and 3639 MEKi-specific peaks (out of 35,114 total MEKi NKX2-1 peaks).

**G.** ChIP-seq tracks showing binding of NKX2-1 to AT1/2 targets in control and MEKi-treated 1027B DP organoids.

**H.** Mean peak profile of control and Trametinib-treated (MEKi) NKX2-1 ChIP-seq signal from DP murine cells (3311) and organoids (1027B) at AT1-specific NKX2-1 binding sites and duodenal HNF4α binding sites.

**FIGURE S7**

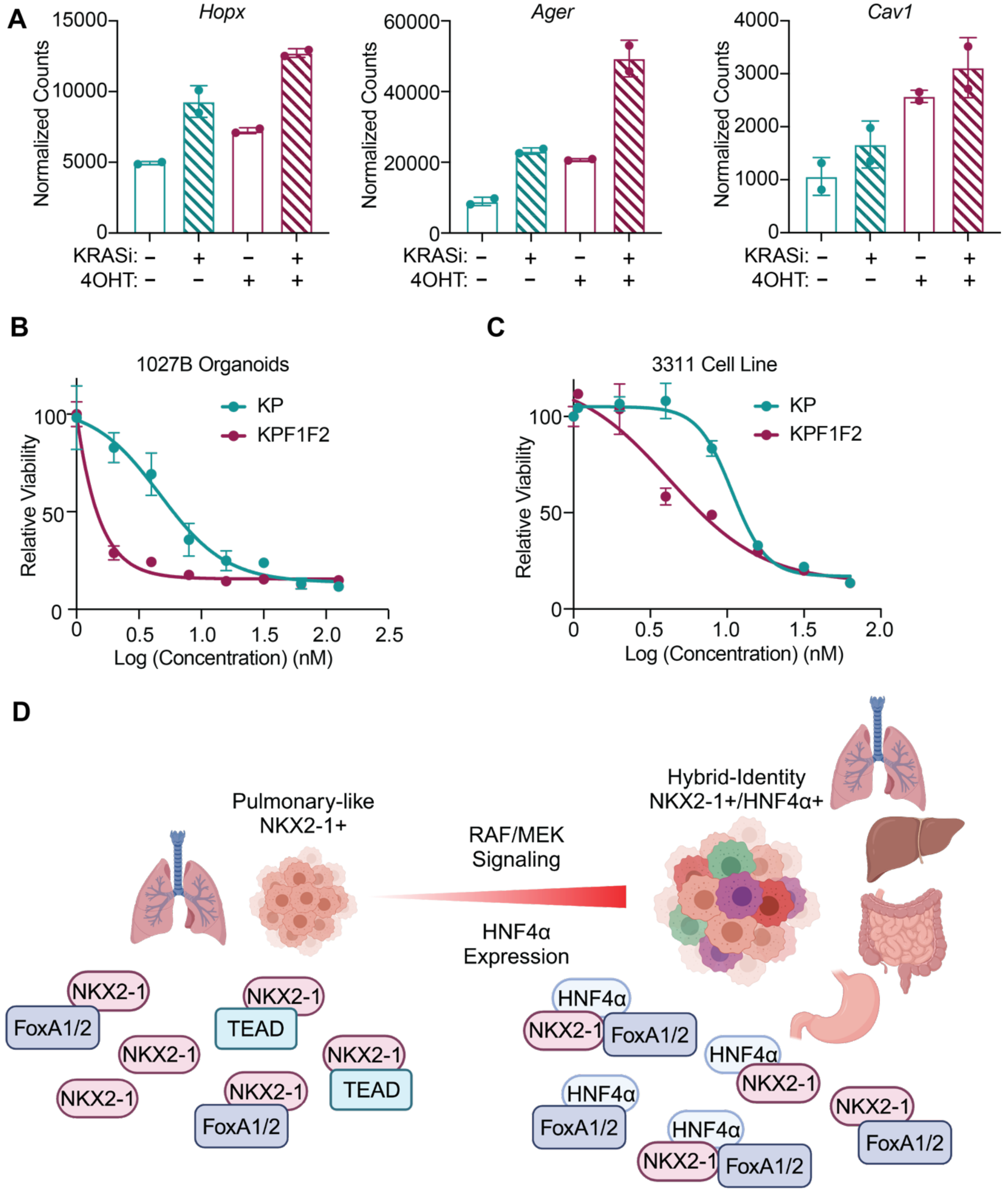
A
Hopx
Normalized Counts
15000
10000
5000
0
KRASi: − + − +
4OHT: − − + +
Ager
Normalized Counts
60000
40000
20000
0
KRASi: − + − +
4OHT: − − + +
Cav1
Normalized Counts
4000
3000
2000
1000
0
KRASi: − + − +
4OHT: − − + +
B
1027B Organoids
KP
KPF1F2
Relative Viability
100
50
0
0 0.5 1.0 1.5 2.0 2.5
Log (Concentration) (nM)
C
3311 Cell Line
KP
KPF1F2
Relative Viability
100
50
0
0 0.5 1.0 1.5 2.0
Log (Concentration) (nM)
D
Pulmonary-like
NKX2-1+
RAF/MEK
Signaling
HNF4α
Expression
Hybrid-Identity
NKX2-1+/HNF4α+
NKX2-1
FoxA1/2
TEAD
HNF4α

**Supplemental Figure 7. Related to Figure 6**.

**A.** Normalized counts from bulk RNA-seq of AT1 marker genes *Hopx, Ager,* and *Cav1* in 3154 DP organoids treated with 3nM MRTX1133 and/or 4OHT to delete *Hnf4a*.

**B-C.** Dose response curves of *Foxa1/2*-conditional control (KP) and deleted (KPF1F2) DP organoid line (**B**) and cell line (**C**) treated with MRTX1133 to inhibit $KRAS^{G12D}$. (**B**) KP $IC_{50}$=4.656nM, KPF1F2 $IC_{50}$=0.059nM. (**C**) KP $IC_{50}$=10.75nM, KPF1F2 $IC_{50}$=4.206nM (one representative replicate of n=2 biological replicates shown).

**D.** Summary of overall model. HNF4α and RAF/MEK signaling induce a pro-growth hybrid-ID state in NKX2-1-positive LUAD.
